# Durable resistance or efficient disease control? Adult Plant Resistance (APR) at the heart of the dilemma

**DOI:** 10.1101/2022.08.30.505787

**Authors:** Loup Rimbaud, Julien Papaïx, Jean-François Rey, Benoît Moury, Luke Barrett, Peter Thrall

## Abstract

Adult plant resistance (APR) is an incomplete and delayed protection of plants against pathogens. At first glance, such resistance should be less efficient than classical major-effect resistance genes, which confer complete resistance from seedling stage, to reduce epidemics. However, by allowing some ‘leaky’ levels of disease, APR genes are predicted to be more durable than major genes because they exert a weaker selection pressure on pathogens towards adaptation to resistance. However, the impact of partial efficiency and delayed mode of action of APR on the evolutionary and epidemiological outcomes of resistance deployment has never been tested.

Using the demogenetic, spatially explicit, temporal, stochastic model *landsepi*, this study is a first attempt to investigate how resistance efficiency, age at the time of resistance activation and target pathogenicity trait jointly impact resistance durability and disease control at the landscape scale. Our numerical experiments explore the deployment of APR in a simulated agricultural landscape, alone or together with a major resistance gene. As a case study, the mathematical model has been parameterised for rust fungi (genus *Puccinia*) of cereal crops, for which extensive data are available.

Our simulations confirm that weak efficiency and delayed activation of APR genes reduce the selection pressure applied on pathogens and their propensity to overcome resistance, but do not confer effective protection. On the other hand, stronger APR genes (which increase selection pressure on the pathogen) may be quickly overcome but have the potential to provide some disease protection in the short-term. This is attributed to strong competition between different pathogen genotypes and the presence of fitness costs of adaptation, especially when APR genes are deployed together with a major resistance gene via crop mixtures or rotations.

## Introduction

In plant pathology, durable resistance and efficient disease control are two important considerations in the use of genetically controlled plant resistance to manage crop diseases (Burdon JJ et al., 2016). Indeed, strategies to deploy plant resistance should first be as efficient as possible to mitigate epidemics and preserve crop health. However, the high evolutionary potential of many plant pathogens means that they can adapt and overcome such resistance, sometimes quickly after deployment in the field (Johnson R, 1983; Parlevliet JE, 2002; García-Arenal F & BA McDonald, 2003). Resistance breakdown results in potentially destructive epidemics and economic losses, leading to increased reliance on pesticides and acceleration of associated environmental issues. In addition, resistance breakdown also means the loss of precious and non-renewable genetic resources, and the need to develop new resistant cultivars, a long and costly process (Zhan J et al., 2015). Therefore, in addition to the provision of efficient crop protection in the short-term, resistance must also be durable, even if these two goals are not necessarily compatible (van den Bosch F & CA Gilligan, 2003; Papaïx J et al., 2018; Rimbaud L et al., 2018a). In this context, simulation models provide powerful tools to explore and evaluate different crop deployment strategies with respect to their epidemiological and evolutionary outcomes, while circumventing the logistical challenges associated with field experiments at large spatio-temporal scales (Rimbaud L et al., 2021).

Plant breeding has typically focused on resistance conferred by major-effect genes, which often confer complete resistance, such that pathogens are unable to infect cultivars carrying those genes. Most major genes encode for an immune receptor of the nucleotide-binding leucine-rich repeat (NLR) protein family, which triggers the immune response (often involving a hypersensitive reaction) after recognition of a pathogen effector (de Ronde D et al., 2014; Gallois J-L et al., 2018). Nevertheless, pathogens may escape this recognition after mutation or suppression of this effector, leading to the restoration of infectivity and resistance breakdown. In these cases, the plant-pathogen genetic interaction is best described by the ‘gene-for-gene’ (GFG) model, according to which the occurrence of disease depends on whether or not the plant carries a resistance gene, and whether or not the pathogen possesses the matching effector (Flor HH, 1955). The scientific literature describes numerous examples of major resistance genes being rapidly overcome by fungi (Johnson R, 1983, 1984; McDonald BA & C Linde, 2002; Parlevliet JE, 2002; Stuthman DD et al., 2007; Park RF, 2008; Burdon JJ & PH Thrall, 2014), bacteria (McDonald BA & C Linde, 2002; Parlevliet JE, 2002), viruses (García-Arenal F & BA McDonald, 2003; Lecoq H et al., 2004; Moury B et al., 2010), and nematodes (McDonald BA & C Linde, 2002), although some of them have maintained effectiveness for many years. Such resistance breakdown results from the high selection pressure experienced by pathogen populations in the presence of such resistance, since only adapted individuals can infect resistant hosts. ‘Resistance-breaking’ mutants may be initially present in the population at low frequency, derive from other pathogen genotypes by mutation or recombination, or be introduced from distant areas through migration. In such cases, the frequency of the mutant genotype increases as it will be strongly favoured by selection and the whole host population may end up infected (Johnson R, 1983, 1984; Lecoq H et al., 2004; Moury B et al., 2010).

Resistance is, however, not always complete or continuous in time. Whether they may be insufficiently expressed, dependent on environmental conditions or simply weak, resistance genes sometimes confer only partial protection to pathogens. In this context, ‘resistance efficiency’ is a key component of partial resistance, and describes how well the infectious cycle of the pathogen is mitigated, i.e., the extent of reduction of one or several pathogenicity traits, such as infection rate, latent or infectious period durations, and reproduction rate (Parlevliet JE, 1979; Lannou C, 2012). Resistance may also be specific to certain host developmental phases (Barrett LG & M Heil, 2012), such as is the case for adult plant resistance (APR, also called ‘mature plant resistance’; Develey-Rivière M-P & E Galiana, 2007). APR genes are often described as being only expressed in adult plants (Burdon JJ et al., 2014; Niks RE et al., 2015), with an efficiency varying from 0% to 100% and depending on plant age and environment (Krattinger SG & B Keller, 2016). However, moderate levels of expression of APR genes can sometimes be already detected in young plants (Park RF & RG Rees, 1989; Cromey MG, 1992; Broers LHM, 1997; Sandoval-Islas JS et al., 2007; Qamar M et al., 2012). This expression tends to increase progressively and the date at which APR genes reach their maximal efficiency (referred to as ‘age of resistance activation’ hereafter) depends on the resistance gene and may occur as late as the anthesis stage (Ma H & RP Singh, 1996). Many APR genes against rust fungi have been documented in cereal crops (page 56 in Burdon JJ, 1987; McIntosh RA et al., 1995; Boyd LA, 2005). They can impact all pathogenicity traits associated with the pathogen infectious cycle: infection rate (e.g. Lr34-Yr18; Qamar M et al., 2012), latent period (Lr16-Lr18, Lr34-Yr18; Tomerlin JR et al., 1983; Elahinia SA & JP Tewari, 2005; Qamar M et al., 2012), sporulation rate (Lr16-Lr18; Tomerlin JR et al., 1983), sporulation duration (Lr16-Lr18; Tomerlin JR et al., 1983). Nonetheless, a wide panoply of molecular mechanisms may underpin APR resistance and these are poorly known (Develey-Rivière M-P & E Galiana, 2007; Krattinger SG & B Keller, 2016). Exceptions include three resistance genes against leaf, stem and yellow rusts of wheat: Lr67 encoding a hexose transporter (Moore JW et al., 2015); Lr34 encoding an ATP-binding cassette (ABC) transporter (Krattinger SG et al., 2009); and Yr36 encoding a chloroplast-localised kinase protein involved in detoxification of reactive oxygen species (Fu D et al., 2009, see also Develey-Rivière M-P & E Galiana, 2007 for resistances against other pathogens).

To the best of our knowledge, the role of delayed activation of plant resistance in disease management and pathogen evolution has never been investigated in simulation models (Rimbaud L et al., 2021), despite its supposed potential to promote resistance durability. Complete resistance is often assumed in modelling studies, and always considered active from the seedling stage. Yet, hosts are thought to generate different selective pressures on pathogens if they express complete, partial or delayed resistance (Stuthman DD et al., 2007; Pilet-Nayel M-L et al., 2017). While complete resistance exerts strong selection on the pathogen to restore infectivity, the pressure imposed by partial and delayed resistances (such as the one conferred by APR genes) is likely lower since they allow some ‘leaky’ levels of disease. One the one hand, partial resistance imposes constant and weak selection pressure on the pathogen; on the other hand, delayed resistance induces a sudden change in the direction of the selection pressure at the time of activation. Both resistances have thus the potential to slow down the speed of pathogen evolution compared to typical major resistance genes. This slower pathogen evolution comes nonetheless at the price of weaker protection against disease, hence the potential of such resistance for disease management is still intriguing, particularly when deployed in conjunction with major gene resistance.

The aim of the present study is to investigate how resistance efficiency, age of resistance activation and target pathogenicity trait of a resistance gene jointly impact resistance durability and epidemiological disease control. Additionally, because deploying different types of resistance is likely a promising approach to benefit from their respective advantages, we also investigate the best strategies to combine a major resistance gene with an APR gene. To study these questions, we use a general simulation framework implemented in the R package *landsepi* (Rimbaud L et al., 2018b). The model is flexible enough to vary parameters related to the deployed resistance genes, and to encompass various pathogen epidemiological traits. Thus, although this work is motivated by rust diseases of cereal crops (for which there is considerable empirical data), our broad conclusions may, to some extent, apply to numerous pathosystems.

## Methods

### Model overview

We used a demogenetic, spatially explicit, temporal and stochastic model developed to explore different plant resistance deployment strategies in agricultural landscapes and evaluate their epidemiological and evolutionary outcomes. A description of the mathematical model is detailed in a previous article (Rimbaud L et al., 2018c) and presented in **Text S1**. Briefly, the model simulates the spread (by wind) and evolution (via mutation) of a clonal spore-borne pathogen in a cropping landscape where susceptible and resistant cultivars are cultivated with controlled proportions and controlled level of spatial aggregation. In the simulated landscape, resistance genes may be deployed in a single host cultivar as a pyramid, or in different cultivars that can be segregated in a mosaic of fields, combined within the same field as mixtures, or alternated within crop rotations. Resistance genes may target one or several pathogenicity traits (reduction of infection rate, sporulation rate or sporulation duration, lengthening of latent period duration) with complete or partial efficiency. The pathogen has the potential to adapt to each of the deployed resistance genes independently, via single mutations (leading to the emergence of new pathogen strains), possibly associated with a fitness cost on cultivars that do not carry the corresponding resistance genes. The spatial unit of the model is the field. The pathogen is disseminated across fields of the landscape using the integral of a power-law dispersal kernel: 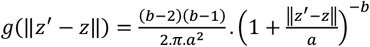 with ‖*z*′ − *z*‖ the Euclidian distance between locations z and z’ in fields i and i’, respectively, a the scale parameter and b a parameter linked to the width of the tail. The plant infection and immune status is modelled using a traditional SEIR (‘susceptible-exposed-infectious-removed’) framework. Plant harvests occur at the end of each cropping season, imposing potential bottlenecks (and thus genetic drift due to randomness in the off-season survival) on the pathogen population. The model was parameterised using available data from the empirical literature to represent wheat rust infection caused by a range of fungal pathogens in the genus *Puccinia* (**Table 1**, details on model calibration in Rimbaud L et al., 2018c, supporting information).

**Table 1.**
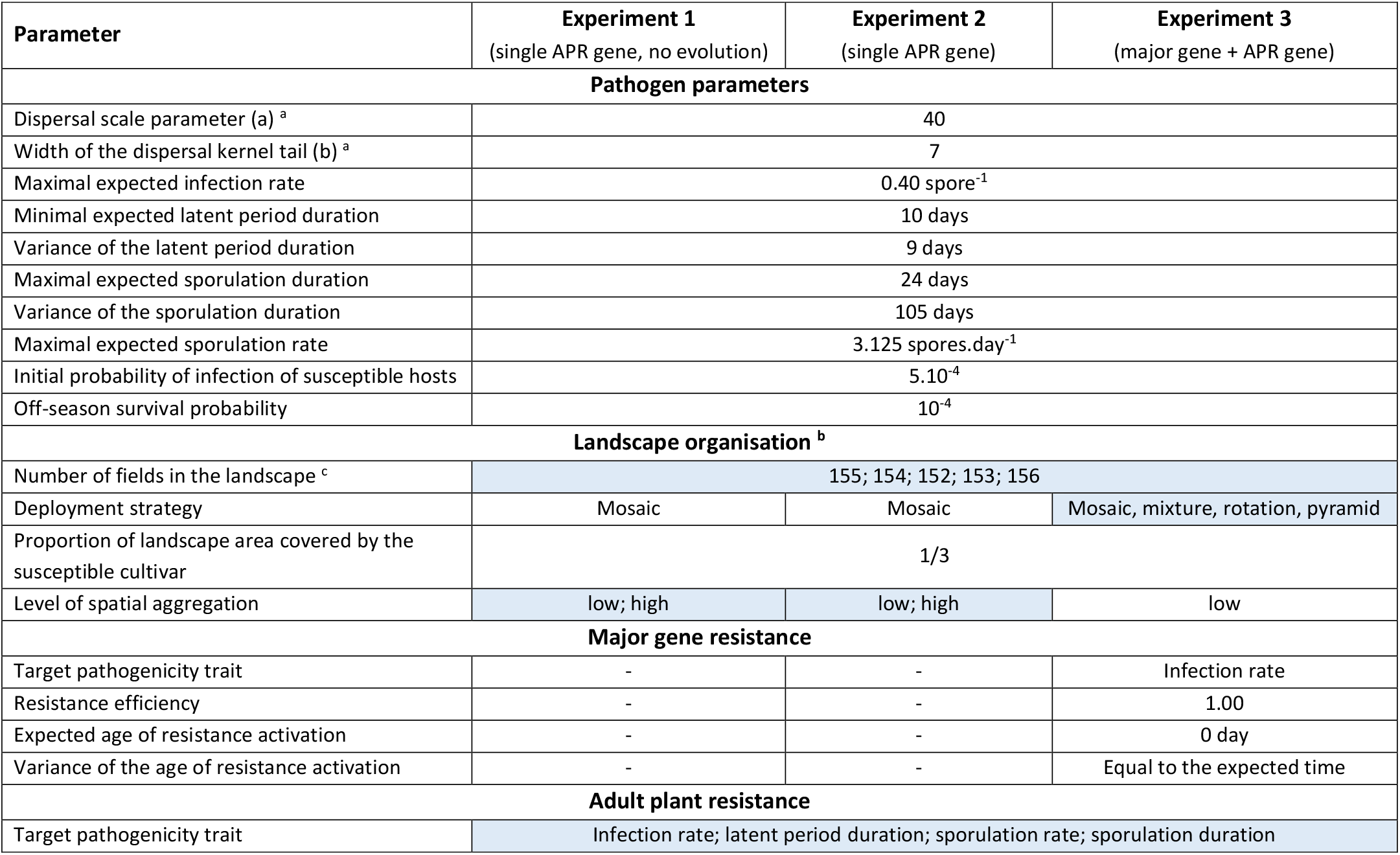
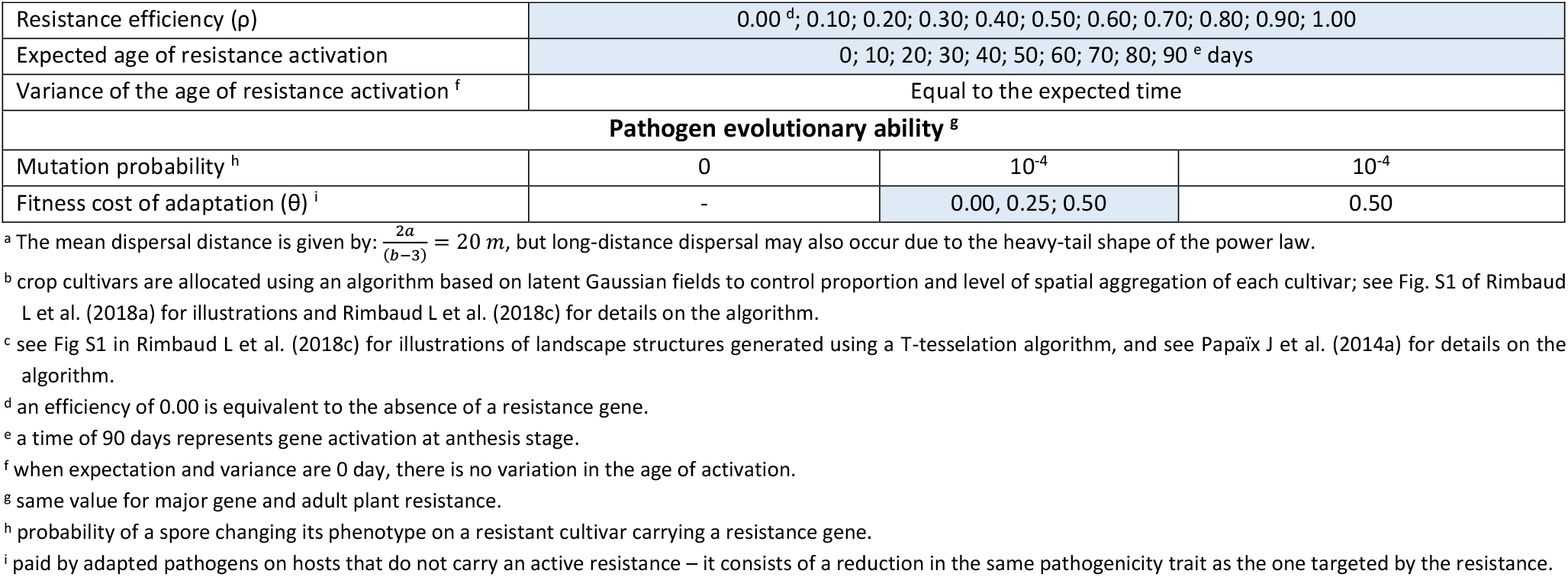
Model parameter and simulation experiments. See Text S1 in Rimbaud L et al. (2018c) for calibration details. Parameters of interest (blue cells) were varied according to a complete factorial design. Every simulation was replicated 10 times x 5 landscape structures to account for stochasticity, resulting in a total of 44,000, 132,000 and 88,000 simulations for the three numerical experiments, respectively.

In this study, the *landsepi* model was extended to include resistance genes with a delayed activation (i.e., APR genes). Cultivars that carry an APR gene are susceptible at the beginning of the cropping season and become resistant once the gene activates. The age of resistance activation is drawn from a gamma distribution every year and for every field planted with a cultivar carrying an APR gene (thus, for a given year, the age of resistance activation is the same for all resistant plants of the same field). For convenience, this distribution is parameterised with the expectation and variance of the age of activation and these are supposed equal (i.e. larger ages are also more variable). The usual shape and scale parameters of a Gamma distribution, β_1_ and β_2_, can be calculated from the expectation and variance, exp and var, with: 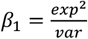 and 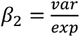, respectively. Both parameters, as well as the target pathogenicity trait and efficiency of resistance, are assumed to be genetically determined and thus characteristic of a given APR gene. Given the parameterisation to wheat rust, if, for example, an APR gene targets the infection rate with an efficiency of 75% and an age of activation of 30 days, the expected infection rate of a healthy resistant host by a spore from a non-adapted (i.e., ‘wild-type’, wt) pathogen will be 0.40 spore^-1^ (see **Table 1**) until resistance activates, after which this rate drops by 75% (i.e., 0.10 spore^-1^) until the end of the cropping season (nothing changes for already infected hosts). If the APR gene targets the latent period duration, hosts infected by a wt pathogen will have an expected latent period of 10 days until resistance activates, after which latent period is increased by 75% (i.e., 17.5 days). Finally, if the APR gene targets the sporulation rate (respectively duration), the expected sporulation rate (resp. duration) will be 3.125 spores.day^-1^ (resp. 24 days) before resistance activation, and 0.781 spores.day^-1^ (resp. 6 days) afterwards.

The model is fully described in **Text S1** and available in the R package *landsepi* version 1.1.1 (Rimbaud L et al., 2018b).

### Numerical experiments

Three successive numerical experiments were carried out to explore APR. Experiment 1 is a baseline scenario destined to evaluate how the deployment of a single APR gene mitigates epidemics in absence of pathogen evolution (i.e., here epidemics are caused by a single pathogen strain, not adapted to the APR gene). Experiment 2 reproduces the same scenario but includes pathogen evolution, to measure the durability of the APR gene and the epidemiological impact of the possible presence of adapted pathogen genotypes. Finally, Experiment 3 investigates whether APR genes and major resistance genes are competing alternatives or can be complementary to each other via appropriate spatio-temporal deployment strategies. **Table 1** summarises model parameters of interest.

In all these experiments, three parameters were systematically allowed to vary: resistance efficiency, age of resistance activation and target pathogenicity trait. Using this approach, we were able to explore a wide range of situations, from the absence of resistance (if resistance efficiency is 0%, **Fig. 1**) to a completely efficient major gene (if efficiency is 100% and there is no delay in resistance activation) with all possible intermediate situations (partially-efficient major gene, completely-efficient APR gene, partially-efficient APR gene).

**Figure 1.**
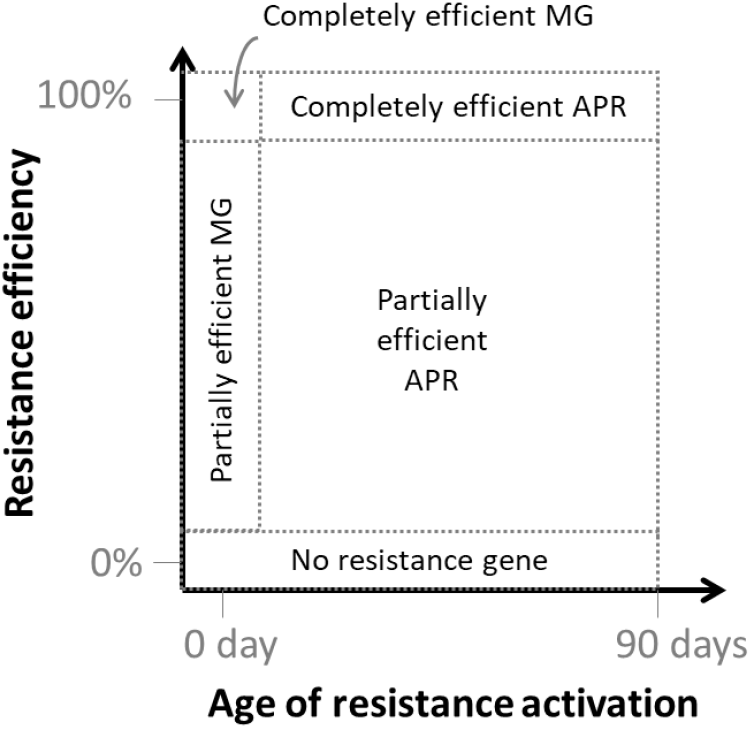
Conceptual exploration of parameters associated with resistance genes: efficiency and age of activation. This formal framework encompasses a wide range of situations. MG: major gene; APR: adult plant resistance.

In the first two experiments, the landscape (representing approximately J=150 fields, total area: 2×2 km^2^, see Fig S1 in Rimbaud L et al., 2018c) was composed of a mosaic of a susceptible (1/3 of total surface) and a resistant cultivar (2/3 of total surface) across the simulated landscape. Cultivars were randomly allocated to fields within the landscape either at low or at high degree of spatial aggregation (**Fig. 2**, left-hand column). The resistant cultivar carried a resistance targeting either infection rate, latent period duration, sporulation rate, or sporulation duration of the pathogen. Analysis of field and greenhouse trials on rust diseases of cereal crops revealed that resistance against these pathogenicity traits measured in different host genotypes can vary from 0% to 100% compared to the most susceptible cultivars (**Table S1**). Thus, in our simulations, resistance efficiency (ρ) was varied from 0 to 100% with increments of 10%. The expected age of resistance activation varied from 0 to 90 days with increments of 10 days; an age of activation of 90 days (the whole epidemic season being 120 days) represents the case where the gene activates at anthesis stage. In the first experiment, the pathogen was not allowed to evolve, whereas in the second, it could adapt to the APR gene through a single mutation. In this case, the impact of fitness cost of adaptation (where fitness cost was defined in terms of loss of pathogenicity on the susceptible cultivar) was studied using three different cost values (θ = 0.00, 0.25, 0.50). Therefore, in this experiment there were two possible pathogen genotypes: the wild-type (‘wt’) and the resistance-breaking (‘rb’) strains, respectively not adapted and adapted to the APR, whose performances on the different cultivars are summarised in **Table 2**.

**Figure 2.**
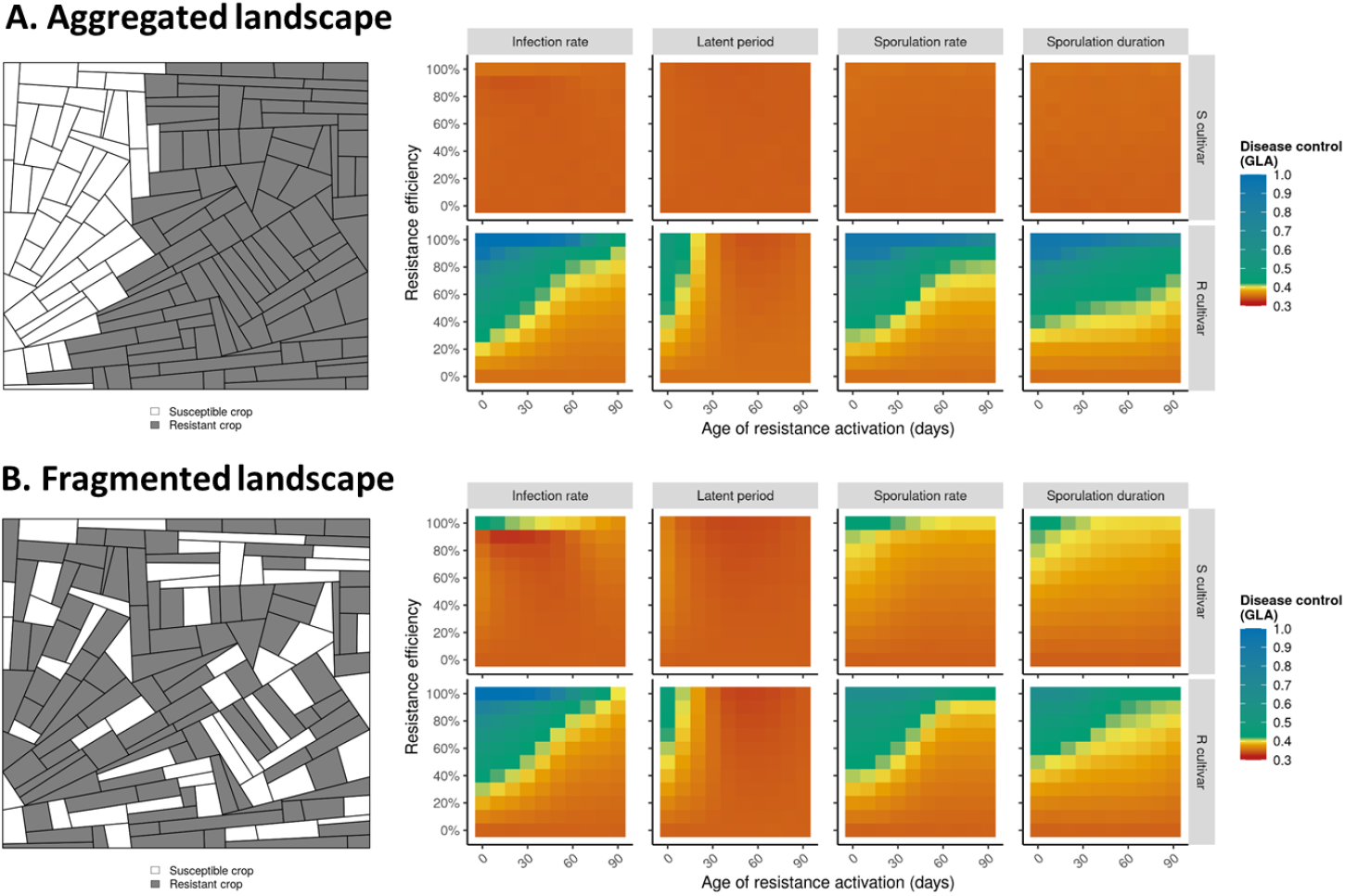
Simulated landscapes (examples on the left) and heatmaps (on the right) of the level of epidemiological control (i.e., disease limitation, measured by the Green Leaf Area, ‘GLA’) in the absence of pathogen evolution for different levels of resistance efficiency (vertical axis), age of resistance activation (horizontal axis) and target pathogenicity traits (columns), for strong (A) or weak levels of spatial aggregation.

**Table 2.** Plant-pathogen interaction matrix with a single resistance gene. The table shows the coefficients by which the value of the target pathogenicity trait (see **Table 1**) is multiplied (except for latent period duration which varies in a direction opposite to that of the other traits: 1-ρ is replaced by 1+ρ and 1-θ is replaced by 1+θ). The coefficients reflect the relative performance of the different pathogen genotypes on the different cultivars. ρ is the efficiency of the resistance gene and θ is the fitness cost of adaptation.

|  | Susceptible cultivar | Resistant cultivar (APR) |  |
| --- | --- | --- | --- |
|  |  | Non-active | Active |
| wild-type pathogen (wt) | 1 | 1 | $1-\rho$ |
| resistance-breaking pathogen (rb) | $1-\theta$ | $1-\theta$ | 1 |

In the third numerical experiment, a major resistance gene and an APR gene were jointly deployed according to one of four strategies: pyramiding, mixture, rotation or mosaic. The major resistance gene was assumed to target pathogen infection rate with complete efficiency and to be active from the beginning of the cropping season. Target pathogenicity trait, resistance efficiency and age of activation of the APR gene were varied exactly as in the first two experiments. In this experiment, there are four possible pathogen genotypes, whose performances on the different cultivars are summarised in **Table 3**.

**Table 3.** Plant-pathogen interaction matrix with two resistance genes, giving the coefficients by which the value of the target pathogenicity trait (see **Table 1**) is multiplied (except for latent period duration which varies in a direction opposite to that of the other traits: 1-ρ is replaced by 1+ρ and 1-θ is replaced by 1+θ). It reflects the relative performance of the wild-type (wt) and the resistance-breaking (rb_1_, rb_2_, rb_12_) pathogen genotypes on the susceptible (S) and resistant cultivars carrying a major resistance gene (MG; cultivar R_1_), an APR gene (R_2_) or both (R_12_). ρ is the efficiency of the APR gene, and θ is the fitness costs of adaptation.

| | S | $R_1$ (MG) | $R_2$ (APR) | | $R_{12}$ (MG+APR) | |
| --- | --- | --- | --- | --- | --- | --- |
|  |  |  | Non-active | Active | Non-active | Active |
| wt | 1 | 0 | 1 | $1-\rho$ | 0 | 0 |
| $rb_1$ | $1-\theta$ | 1 | $1-\theta$ | $1-\rho$ | 1 | $1-\rho$ |
| $rb_2$ | $1-\theta$ | 0 | $1-\theta$ | 1 | 0 | 0 |
| $rb_{12}$ | $(1-\theta)^2$ | $1-\theta$ | $(1-\theta)^2$ | $1-\theta$ | $1-\theta$ | 1 |

For this experiment, spatial aggregation was fixed at a low value (representing a fragmented landscape), and the fitness cost of pathogen adaptation to θ=0.50. Indeed, results obtained in the second experiment showed that this parameterisation maximises the interaction between cultivars (in terms of pathogen dispersal and competition between pathogen genotypes) within a spatial deployment strategy. For all deployment strategies, 1/3 of the landscape was composed of the susceptible cultivar. The remaining 2/3 were occupied either by a single cultivar carrying the two genes (pyramid strategy), a mixture (in every field) of two resistant cultivars in balanced proportions (each cultivar carrying one of the two genes; mixture strategy), a rotation of these two resistant cultivars (every year; rotation strategy), or a mosaic of the two resistant cultivars in balanced proportions (every cultivar representing 1/3 of the landscape area; mosaic strategy) (**Fig. S6**).

Simulations were run for T=120 time-steps per cropping season over a time period of Y=30 years. Initially, only the wild-type pathogen (i.e., not adapted to any resistance), ‘wt’, was present in susceptible hosts, with a probability of any host being initially infected of 5.10^−4^. The wt strain is unable to infect resistant hosts carrying an APR gene only if resistance is both complete and active. In all other situations, the wt strain is able to infect the hosts carrying an APR gene. In any case, a single mutation (with probability 10^−4^, except in the first experiment where evolution did not occur) is required to overcome a resistance gene (should it be a major gene or an APR gene) and restore complete pathogenicity, in conformity with a gene-for-gene interaction. Thus, two distinct mutations are required to generate the rb_12_ genotype from the wt genotype, and overcome the pyramid composed of a major resistance gene and an APR gene.

Model stochasticity includes field shape and boundaries, cultivar allocation to the different fields within the simulated landscape, age of APR gene activation, pathogen dispersal, mutation, off-season survival, and SEIR transitions. To account for this stochasticity, simulations were run on five different landscape structures and replicated 10 times, resulting in 50 replicates for every parameter combination. Thus, the complete factorial design of the three experiments resulted in a total of 44,000, 132,000 and 88,000 simulations, respectively.

### Model outputs

In this work, epidemiological control is defined as the ability of a given deployment strategy to reduce disease impact on the resistant cultivar(s). Here, it is measured by the relative green leaf area (GLA), i.e., the proportion of healthy hosts relative to the total number of hosts, averaged for every cultivar across the whole simulation run. The higher the value of the GLA, the better the epidemiological control. Let’s say that, in accordance with the SEIR architecture of the model, H_i,v,t_, L_i,v,p,t_, I_i,v,p,t_ and R_i,v,p,t_, respectively denote the number of healthy, latent, infectious and removed host individuals in field i (i=1,…,J), for cultivar v (v=1,…,V), pathogen genotype p (p=1,…,P) at time-step t (t=1,…,TxY). Then, the GLA of cultivar v can be computed as follows:

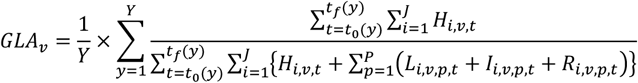

Evolutionary control is quantified here using resistance durability (for experiments 2 and 3), which measures the ability of a given deployment strategy to limit pathogen evolution and delay resistance breakdown (i.e., emergence of the ‘rb’ strain). Durability is evaluated using the time t* when the number of resistant hosts infected by the rb strain (e.g., if v=2 for the resistant cultivar and p=2 for the rb pathogen, 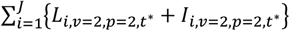) exceeds a threshold above which extinction of this strain is unlikely (fixed at 50,000, see Rimbaud L et al., 2018c, supporting Text S2 for details). To understand the contribution of the different pathogen genotypes to an epidemic, we also calculate, across the whole simulation run and for every cultivar, the proportion of infections due to each pathogen genotype relative to all infections. The contribution of pathogen p to epidemics on cultivar v is computed as follows:

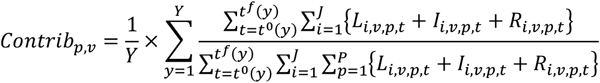

## Results

Three separate numerical experiments were carried out to investigate the epidemiological and evolutionary outcomes of deployment strategies based on APR: the first two experiments were performed with an APR gene alone, and the third with a combination of an APR gene and a major resistance gene.

### Experiment 1: Deployment of a single APR gene in a susceptible landscape with no pathogen evolution

Disease control, measured by the Green Leaf Area averaged for every cultivar across the whole simulation run, was first evaluated when a single APR is deployed in the landscape and the pathogen does not have the possibility to overcome the resistant cultivar.

As expected, for the resistant cultivar, disease control increases with higher efficiency and smaller age of resistance activation (**Fig. 2**). Globally, the target pathogenicity trait offering the best level of disease control is the infection rate when resistance is activated early in the cropping season, whereas it is the sporulation duration when resistance is activated late (**Figs. 2 & S1**). On the susceptible cultivar, disease control is globally poor except when the level of spatial aggregation between cultivars is low and the APR carried by the resistant cultivar is almost completely efficient, activates very early (i.e., it is roughly similar to a major gene), and targets the pathogen infection rate, sporulation rate or sporulation duration (**Fig. 2B**). This comes at the price of a slightly decreased level of control for the resistant cultivar compared to an aggregated landscape.

### Experiment 2: Deployment of a single APR gene in a susceptible landscape with pathogen evolution

In this experiment, the pathogen can adapt to the APR gene via mutation, leading to the emergence of a rb strain.

#### Impact of resistance efficiency and age of activation

Regardless of the target pathogenicity trait, fitness cost and level of spatial aggregation, the results indicate that weak resistance (whether it is inefficient or delayed in activation; bottom right corner of graphics in **Figs. 3, S2, S3, S4**) is always durable (**panels A and B**), meaning that rb pathogen genotypes never emerged in the 30-year simulations (**panels E and F**). However, in this situation, resistance does not confer good epidemiological protection against the wt pathogen, as shown by the second output variable (‘Disease control’, **panels C and D**). In contrast, strong resistance (highly efficient and activated early in the growing season; top left corner of graphics in **Figs. 3, S2, S3, S4**) shows poor durability (**panels A and B**), indicating that the rb pathogen genotype quickly emerged and invaded the resistant host population (**panels E and F**). This again results in poor epidemiological control for the resistant cultivar (**panels C and D**). However, when fitness costs are large (θ=0.50), there is a critical zone (i.e., a range of parameter values leading to optimal epidemiological control) where disease control by the resistant cultivar reaches a higher level, particularly when infection rate is targeted by the APR gene. This zone corresponds to resistance efficiencies higher than 60% and age of activation between roughly 30 and 80 days (**Fig. 3CD**). **Fig. S5** illustrates examples of simulations carried out in the three contrasted scenarios described just above (weak resistance, strong resistance, critical zone).

**Figure 3.**
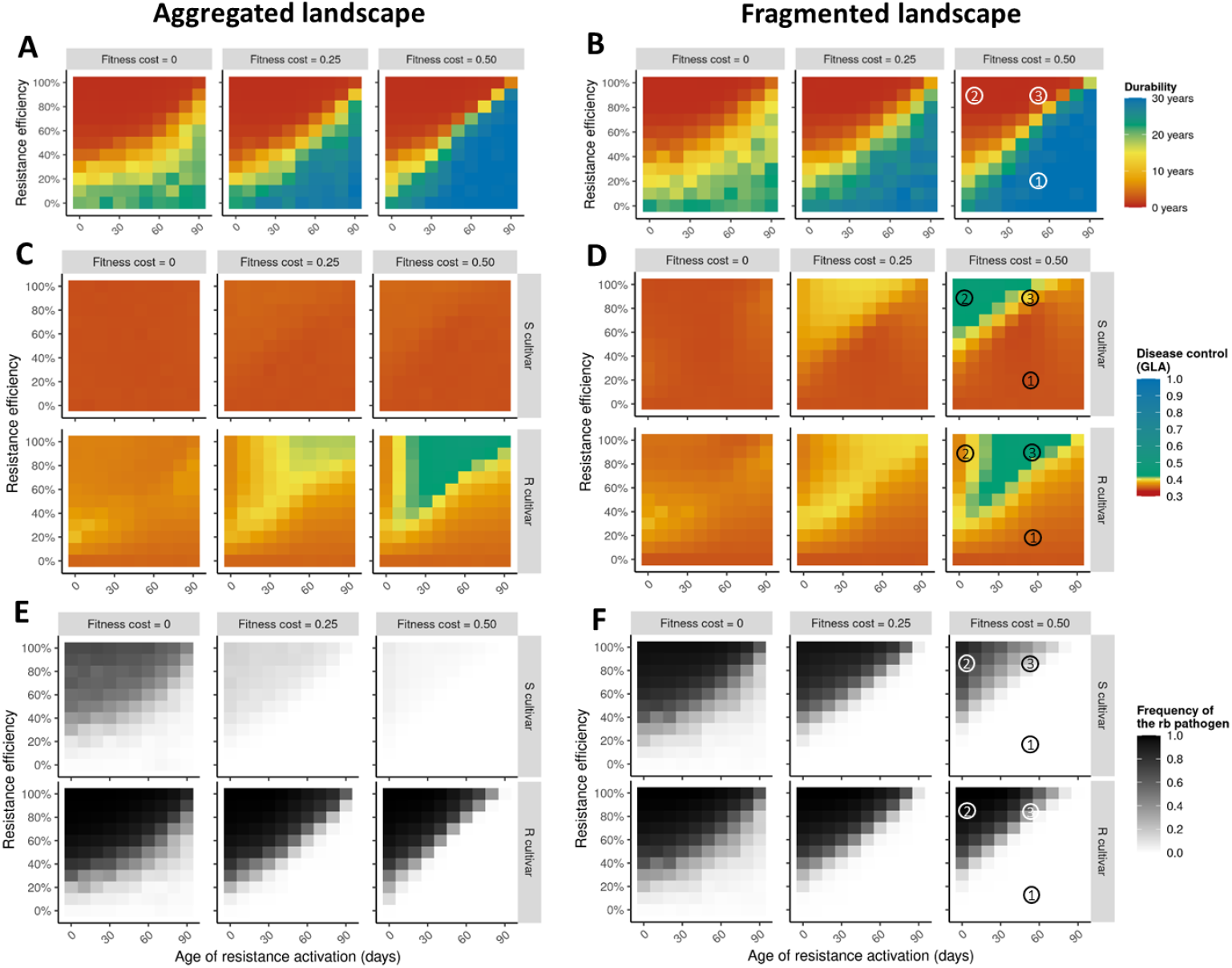
Heatmaps of the levels of evolutionary control (resistance durability as measured by the number of years before the emergence of the resistance-breaking (‘rb’) pathogen genotype; panels A and B), epidemiological control (i.e., disease limitation, measured by the Green Leaf Area (‘GLA’) on the susceptible (‘S’) and the resistant (‘R’) cultivars; panels C and D) and average frequency of the rb pathogen (panels E and F) for different levels of resistance efficiency (vertical axis), age of resistance activation (horizontal axis) and fitness cost of pathogen adaptation (columns), for strong (panels A, C, E) or weak (B, D, F) levels of spatial aggregation. The target pathogenicity trait is the infection rate. Circled numbers refer to example simulations in **Fig. S5**.

#### Impact of fitness cost of adaptation

Decreasing the loss of pathogenicity of the rb pathogen on the susceptible cultivar (effect of columns in **Figs. 3, S2, S3, S4**) tends to decrease both durability and disease control (at intermediate resistance efficiency and delay of activation, rb genotypes emerge more often and cause more damage). In particular, when there are no fitness costs of adaptation, the critical zone previously described disappears completely.

#### Impact of the level of field spatial aggregation

The strongest impact of spatial aggregation is on the genetic composition of the pathogen population and the associated epidemic damage (**Figs. 3, S2, S3, S4, panels C to F**). The susceptible cultivar is mostly infected by the wt pathogen in aggregated landscapes, leading to severe epidemics. In contrast, in fragmented landscapes, for strong resistance (highly efficient or activated early in the growing season) and in presence of fitness costs of adaptation, the susceptible cultivar is mostly infected by the rb pathogen, resulting in moderate to good epidemiological control (due to the fitness penalty). Conversely, epidemiological control for the resistant cultivar seems slightly better in aggregated landscapes (especially when resistance is strong but considerably delayed in the cropping season, top right corner of heatmaps, **Fig. 3CD**). In the absence of fitness costs of adaptation or for weak resistance (inefficient or activated late in the growing season), the genetic composition of the pathogen is similar on the two cultivars, and the associated damage is high.

#### Impact of the target pathogenicity trait

All the previous results hold qualitatively with the different pathogenicity traits targeted by resistance. When resistance targets sporulation rate or the duration of the sporulation period, the genetic composition of the pathogen population and the level of evolutionary control (resistance durability) are similar to what was observed for the infection rate (**Figs. S3, S4**). There are, however, quantitative changes in the epidemiological outcome, as size and location of the critical zone are slightly different depending on the target pathogenicity trait. For infection rate, as mentioned before, the critical zone of good disease control corresponds to resistance efficiencies higher than 60% and activation between 30 and 80 days. For sporulation rate (or sporulation duration), the critical zone corresponds to efficiencies higher than 80% (respectively 90%) and activation after 50 days (respectively 80 days). Resistances increasing the duration of the latent period and having a high efficiency and a delayed activation (more than 30 days, **Fig. S2**, top right corner of graphics) are more durable than those targeting the other traits. This is a consequence of the absence of emergence of the rb pathogen. However, the level of epidemiological control is poor in comparison to the other target traits, and the size of the critical zone is considerably reduced (restricted to resistance efficiencies between 80 and 100% and times to activation of less than 20 days).

### Experiment 3: Simultaneous deployment of a major resistance gene and an APR gene in a susceptible landscape

In a third numerical experiment, resistance durability and disease control were evaluated when a major resistance gene and an APR gene were simultaneously deployed across a landscape, either within the same cultivar (R_12_, pyramiding strategy) or in two distinct cultivars (R_1_ and R_2_, respectively) which could be cultivated in different fields (mosaic strategy), within the same field as mixtures, or alternated in time through crop rotations (see **Fig. S6** for examples of simulated landscapes).

#### Impact of resistance efficiency, age of activation and deployment strategy

Regardless of the characteristics of the APR gene (efficiency, age of activation, target pathogenicity trait), the major gene is always overcome quickly after deployment (**Figs. 4, S7, S8, S9**), except when it is pyramided with a very efficient APR gene that is activated early in the growing season (which is essentially the same as a pyramid of two major resistance genes). This rapid breakdown is mostly attributed to the emergence of the single mutant ‘rb1’ (except when the major gene is pyramided with a strong APR, in which case the breakdown is due to the double mutant ‘rb12’, **Fig. 5**). With respect to the durability of the APR gene and the level of protection it confers on the associated cultivar (R2), weak resistance (i.e., inefficient or delayed in activation) is durable (neither the ‘rb2’ nor the ‘rb12’ genotypes emerged) but offers poor protection against the ‘wt’ and ‘rb1’ genotypes (**Fig. 4 & 5**), similar to the results for Experiment 2. When resistance is strong (very efficient and activated early), it is quickly overcome (**Fig. 4**), either by ‘rb2’ in mosaics and mixtures, or by ‘rb12’ in rotations and pyramids (**Fig. 5**). In mosaics, this leads to the same critical zone previously described for Experiment 2. In contrast, in mixtures and rotations, the level of control stays high for a large range of resistance efficiencies and times to activation. In pyramids, there is a good level of control only for highly efficient resistances (**Fig. 4**). For the resistant cultivar carrying the major gene (R1), disease control shows contrasting results depending on the deployment strategy. It is globally poor in mosaics and globally good in rotations. In mixtures, it is good only when the second resistant cultivar (R2) carries a strong APR gene that is activated early. In pyramids, it is good as long as the APR has a strong efficiency. For the susceptible cultivar, a good level of disease control can be obtained if the APR (deployed in cultivar R2) has a strong efficiency and early activation, especially if pyramided with a major gene. In this situation the susceptible cultivar is invaded by both the ‘wt’ and the ‘rb12’ pathogen genotypes (**Fig. 5**).

**Figure 4.**
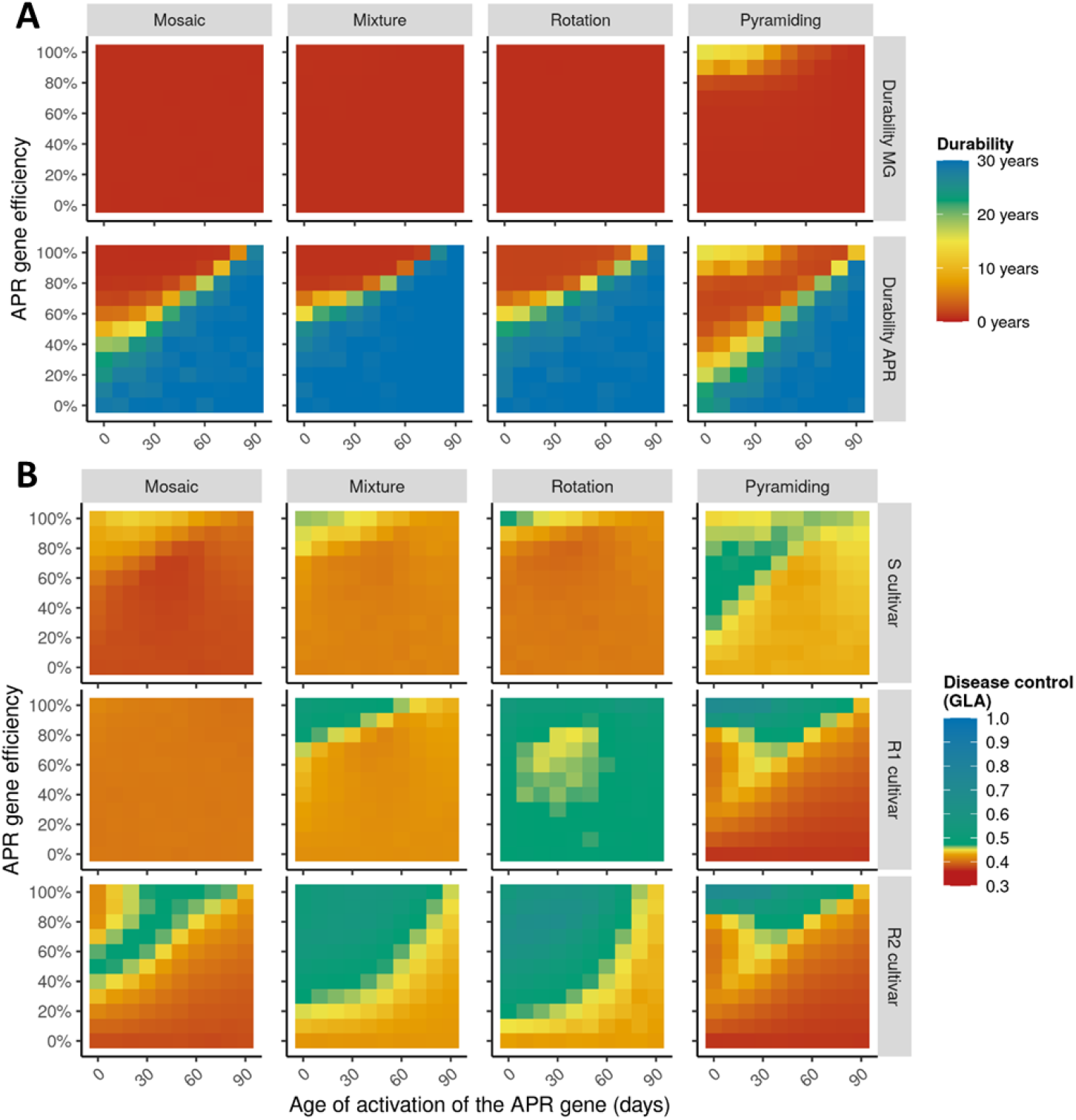
Heatmaps showing the levels of A) evolutionary control (resistance durability, measured by the number of years before the emergence of resistance-breaking genotypes) and B) epidemiological control (i.e., disease limitation, measured by the Green Leaf Area, ‘GLA’) on a susceptible cultivar ‘S’, a resistant cultivar ‘R1’ carrying a completely efficient major gene (‘MG’) and a resistant cultivar ‘R2’ carrying an APR gene, for different levels of APR efficiency (vertical axis), age of APR activation (horizontal axis) and deployment strategies (columns; note that for pyramiding, R1 and R2 refer to the same cultivar). The target pathogenicity trait of the APR gene is the infection rate, the level of spatial aggregation is low, and the fitness cost is 0.50.

**Figure 5.**
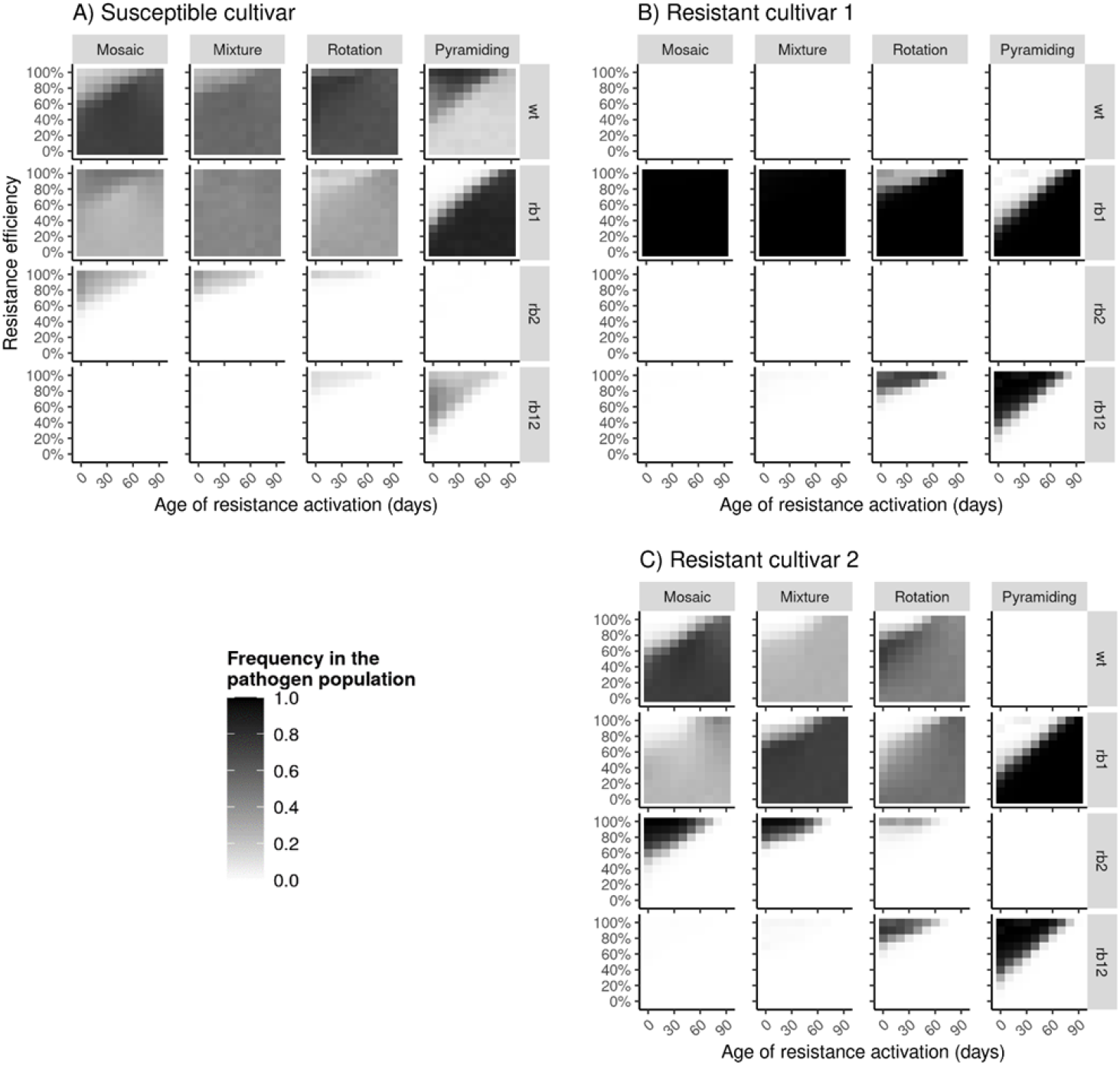
Average frequency of the different pathogen genotypes (see **Table 3** for notations) on a susceptible cultivar ‘S’, a resistant cultivar ‘R1’ carrying a completely efficient major gene and a resistant cultivar ‘R2’ carrying an APR gene, for different levels of APR efficiency (vertical axis), age of APR activation (horizontal axis) and deployment strategies (columns; note that for pyramiding, R1 and R2 refer to the same cultivar). The target pathogenicity trait of the APR gene is the infection rate, the level of spatial aggregation is low, and fitness cost is 0.50.

#### Impact of targeted pathogenicity trait

The results are qualitatively the same when sporulation rate and sporulation duration are targeted by the APR gene instead of the infection rate (**Figs. S8 & S9**). When resistance conferred by the APR gene increases the length of the latent period (**Fig. S7**), it is durable for a larger range of parameter values (i.e., resistance efficiency and age of activation) compared with the other target traits. However, in this situation the level of epidemiological control for the different cultivars is poor in comparison to the other target traits.

## Discussion

To the best of our knowledge, adult plant resistance (APR) has never been explored in mathematical models dealing with plant resistance deployment (Rimbaud L et al., 2021), despite its presence in numerous resistant cultivars of cereals and other crops (page 56 in Burdon JJ, 1987; McIntosh RA et al., 1995; Boyd LA, 2005; Chen XM, 2005; Develey-Rivière M-P & E Galiana, 2007; Chen W et al., 2014). Therefore, and because APR may affect different pathogenicity traits, in a delayed and potentially incomplete manner, we used the mathematical model implemented in the R package *landsepi* (Rimbaud L et al., 2018c) to explore three parameters associated with this type of resistance: target pathogenicity trait, efficiency and age of activation. The main objective was to evaluate the impact of these parameters on resistance durability (evolutionary pathogen control) and disease limitation (epidemiological control). We designed numerical experiments to explore three scenarios: the deployment of a single APR gene in a susceptible landscape, firstly without and secondly with pathogen evolution. The third experiment assessed the deployment of an APR gene together with a major resistance gene according to different spatiotemporal deployment strategies (**Table 1**).

### Favouring competition offers good epidemiological control in spite of pathogen adaptation

Globally, our results show that an APR gene is never overcome when it is inefficient with respect to reducing the target pathogenicity trait or is activated late in the cropping season (**Figs. 3AB, 4A**). This is due to the weak selection pressure applied to the pathogen population, given that the wt genotype can thrive on cultivars carrying such resistance genes almost as if they were susceptible. This is in accordance with results obtained via different simulation models (Carolan K et al., 2017; Crété R et al., 2020) and confirms one of the mechanisms according to which partially efficient resistance is generally predicted to be more durable than complete resistance (Lecoq H et al., 2004; Stuthman DD et al., 2007; Zhan J et al., 2015). Such phenomena have also been described for pest adaptation to chemicals, where small application doses were shown to slow down the emergence of adapted genotypes (Hobbelen PHF et al., 2014). Partial resistance with low efficiency or delayed activation, however, results in a weak level of epidemiological control (**Figs. 2, 3CD, 4B**). In contrast, when resistance strongly reduces the target pathogenicity trait of the wt pathogen, particularly when this happens early in the cropping season, it has a high potential to protect the resistant cultivar (Experiment 1, **Fig. 2**), as expected in absence of pathogen evolution and already shown in demographic models (e.g., Papaïx J et al., 2014b). However, if pathogen evolution is possible, the high selection pressure leads to the rapid emergence of a rb pathogen which invades the resistant host population, resulting in both low durability and disease control (Experiment 2, **Fig. 3**). This is similar to a scenario where a single major gene (i.e., complete resistance) is deployed in the landscape and quickly overcome (Rimbaud L et al., 2018c).

There is, however, an intermediate region of the parameter space where the APR gene is broken down but still confers a good level of epidemiological protection. This occurs in presence of pathogen evolution only (i.e., in Experiment 2 but not in Experiment 1), and mostly when resistance is delayed in the cropping season but has sufficiently high efficiency once activated. The delay in resistance activation allows the wt genotype to infect resistant hosts early in the season, more efficiently than potential rb genotypes which suffer a fitness cost while resistance is inactive. As soon as it activates, resistance is strong enough to select for rb genotypes, but many hosts are, at this time, already infected by the wt genotype. The ensuing strong competition between the wt and rb genotypes (Experiment 2, **Fig. 3 & S5**) explains the limitation on epidemic development (Keesing F et al., 2006). In this context, a resistant crop carrying an APR may conceptually be seen as a within-season rotation between a susceptible and a resistant cultivar. There is also an interesting parallel to make with induced resistance. In this case, a resistant cultivar becomes, for a while, less susceptible to a rb pathogen when previously primed (but not infected) by a wt pathogen (Calonnec A et al., 1996; Clin P et al., 2021), whereas a cultivar carrying an APR gene cannot be infected by a rb pathogen when previously infected (at the time when resistance was still inactive) by a wt pathogen. Another type of competition occurs when the APR gene has a small efficiency but is activated very early, corresponding to the bottom left part of the critical zone. In this case, the wt genotype is (slightly) limited on the resistant cultivar (due to early resistance activation), while the rb genotype is (slightly) limited on the susceptible cultivar (due to the fitness cost of adaptation). Provided that the pathogen can disperse from on type of cultivar to another, this situation results in a competition between two specialist genotypes for a limited resource (Mikaberidze A et al., 2015; Clin P et al., 2022).

The size of the critical zone (i.e., the range of parameter values leading to optimal epidemiological control for the resistant cultivar) is amplified whenever such competition between pathogen genotypes is stimulated. In our results, this is the case for high fitness costs of pathogen adaptation, which increases the penalty for rb genotypes on both susceptible hosts and hosts with still inactive APR gene and reduces the performance of these genotypes relative to the wt genotype. This corroborates other modelling studies showing that high fitness costs decrease epidemic severity (Pietravalle S et al., 2006; Djidjou-Demasse R et al., 2017; Rimbaud L et al., 2018a; Watkinson-Powell B et al., 2020). In the absence of pathogen adaptation (i.e., when there is only one pathogen genotype, Experiment 1, **Fig. 2**) or fitness cost (Experiment 2, **Fig. 3**) this effect completely disappears.

### The level of spatial aggregation of the landscape impacts interactions between cultivars

A high level of spatial aggregation between cultivars in the landscape (e.g. **Fig. 2A**) isolates cultivars and the respective pathogen genotypes that infect them. In terms of epidemiological control of a susceptible cultivar, it results in severe epidemics caused by the wt pathogen genotype (**Figs. 2A, 3C**). Conversely, in a fragmented landscape (weak level of aggregation, e.g. **Fig. 2B**), the increased connectivity between different cultivars favours pathogen migration from one cultivar to another (Taylor PD et al., 1993). This reduces epidemics on the susceptible cultivar as a result of two different mechanisms which our two first experiments help disentangle. First, there is a dilution effect (Mundt CC, 2002; Rimbaud L et al., submitted), especially in the presence of a cultivar carrying a very strong resistance activated early in the cropping season. Indeed, in this situation, spread of the wt genotype across susceptible fields is mitigated by the intervening presence of resistant hosts (Experiment 1, **Fig. 2B**). This is similar to non-host plants that act as propagule sinks and thus decrease epidemic spread on susceptible plants (Suzuki SU & A Sasaki, 2011; Papaïx J et al., 2014b). Second, competition occurs between different pathogen genotypes when the resistant cultivar has an intermediate to strong efficiency and a delayed activation (as described previously). In this case, rb genotypes emerging in resistant fields disperse to susceptible fields (Experiment 2, **Fig. 3D**). This leads to a reduction in the damage caused to the susceptible cultivar (provided that rb genotypes suffer a fitness cost compared to the wt genotype) (Watkinson-Powell B et al., 2020). The side-effect of such a protective effect of the susceptible cultivar by the resistant cultivar is a slightly reduced level of disease control on the resistant cultivar when resistance is activated late in the cropping season because it is more exposed to wt pathogen genotypes emerging from susceptible fields. Indeed, wt genotypes are fitter than rb genotypes on the resistant cultivar as long as resistance is inactive, due to the presence of fitness costs. Spatial aggregation has been previously demonstrated to have an ambivalent effect on disease management. In fact, earlier modelling work showed that fragmented landscapes better mitigate epidemics on susceptible crops but are more prone to resistance breakdown, compared to aggregated landscapes (Papaïx J et al., 2018; Rimbaud L et al., 2018a).

### Optimal efficiency and age of activation of APR genes depend on the target pathogenicity trait

A recent opinion published by Saubin M et al. (2022) states that life history traits targeted by resistance influences its durability. In fact, in the present work, the size and location of the critical zone in parameter space depends on the pathogenicity trait targeted by the APR gene. When sporulation rate or duration of the sporulation period are targeted, the critical zone is shifted towards higher resistance efficiencies and longer times to resistance activation compared to the situation where infection rate is targeted (top right of **Figs. 3CD, S3CD, S4CD**). This shift occurs probably because sporulation takes place later than infection in the pathogen infectious cycle. Therefore, more time is required for the wt pathogen genotype to generate sporulating lesions in the resistant host population before resistance activation (which will favour rb genotypes). APR genes targeting the latent period duration seem very durable, but offer poor disease control in comparison to APR genes targeting other traits (**Fig. S2**). This is because even when resistance is fully efficient (i.e., latent period is multiplied by 2), pathogen spread is still possible, which imposes weak selection pressure in favour of rb genotype but provides weak protection against the wt pathogen. This conclusion contrasts with published literature suggesting that latent period duration should be the most influent component of pathogen aggressiveness because it determines the number of possible infection cycles on a crop (Parlevliet JE, 1979; Leonard KJ & CC Mundt, 1984; Sandoval-Islas JS et al., 2007). Nevertheless, sensitivity analyses of models simulating epidemics of wheat leaf rust (Kulkarni RN et al., 1982) and potato late blight (Van Oijen M, 1992) have shown that latent period duration was equally or even less influential on disease spread and severity than other pathogenicity traits. These contrasted results highlight the crucial importance of the width of parameter variation ranges in numerical experiments. In our work, the range of variation for resistance efficiency was based on available data for rust fungi. Analysis of the minimal and maximal possible values of the pathogenicity traits measured on different cultivars of cereal crops (**Table S1**) showed that these traits may vary from about 0% to -100% (0% to +100% for latent period duration) relative to the most susceptible cultivar (except sporulation duration, for which there is little data). We thus allowed resistance efficiency to vary from 0 to 100% for all pathogenicity traits.

### Major resistance genes and APR genes can be combined at landscape scale

The deployment of a single major resistance gene in a landscape results in rapid breakdown by the corresponding rb1 pathogen and severe epidemics on both susceptible and resistant cultivars (the bottom line of heatmaps in **Fig. 4** shows the situation where the APR is absent, its efficiency being 0%). Combining a major gene with an APR gene in the landscape generally does not prevent the major gene from being overcome, however it may have interesting synergies in terms of epidemiological control depending on the deployment strategy (**Fig. 4**). As discussed earlier, one of the greatest benefits of APR genes is the limitation of epidemics due to competition between pathogen genotypes. Therefore, the presence of different sources of resistance in the landscape, should they be overcome, increases the number of pathogen genotypes present and thus the number of competitors. Globally, this decreases epidemic damage on all cultivars (Mikaberidze A et al., 2015; Clin P et al., 2022).

More specifically, when a cultivar carrying a major gene is planted in mixtures (i.e., in the same field) with a cultivar carrying an APR gene, the first cultivar benefits from a dilution effect (since only rb1 genotypes can infect it) conferred by the presence of the second one, which itself benefits from strong competition between the wt, rb1 and rb2 genotypes. While to some extent this should also occur in mosaics (i.e., different cultivars segregated in different fields), our results do not show such synergies for the mosaic strategy. This is probably because of the model assumption that the pathogen was initially present in all susceptible fields of the landscape, added to the fact that pathogen dispersal is mostly at the intra-field scale in our parameterisation (**Table 1**). The impact of landscape heterogeneity on epidemic spread via competition and dilution effects might be stronger for pathogens with different life histories (Mundt CC, 2002). Here, the best epidemiological control is obtained when crop cultivars are mixed at the finest spatial grain. Indeed, optimal disease control requires that the spatial scale of disease management matches the scale of pathogen dispersal (Gilligan CA, 2008). When the two resistant cultivars are rotated over time (rotation strategy), pathogen genotypes are confronted by an alternation of strong selection towards the rb1 genotype (when the cultivar carrying the major gene is cultivated) and strong or weak selection towards the rb2 genotype (when the cultivar carrying the APR gene is cultivated). If the APR is not too strong or has a delayed activation, selection towards rb2 is weak, which allows competition between wt, rb1 and rb2 genotypes and reduces epidemics. Otherwise, selection is strong and the genotype that performs best in the system is the double mutant rb12 (generalist genotype able to infect all cultivars). However, this genotype is penalised by severe fitness costs (**Table 3**), which reduces epidemic damage as well. This is in line with a previous modelling study comparing mosaics, mixtures, rotation and pyramids of major resistance genes: rotation had the best epidemiological outcome once all resistances had been overcome (i.e., in the presence of rb genotypes) (Rimbaud L et al., 2018a). Finally, if the major gene and the APR gene are pyramided in the same cultivar and the efficiency of the APR gene is strong enough, the delayed action of the APR gene triggers competition between the single mutant rb1, selected for as long as the APR is inactive, and the double mutant rb12, selected for as soon as the APR activates. This competition reduces epidemic damage on the pyramid cultivar. However, the presence of the APR gene does not prevent the major gene from being overcome, unless it is activated very early in the cropping season. This is in agreement with previous modelling results: durability of a major gene was greater when pyramided with a quantitative resistance (activated from the beginning of the cropping season), but only if the latter exhibited strong efficiency (Rimbaud L et al., 2018c).

### General conclusions, limits and perspectives

There are several nonexclusive arguments for why APR genes are thought to be more durable than traditional major genes. Firstly, it could be inherent to the molecular mechanism of APR genes, that may be more difficult for the pathogen to overcome than classical NLR proteins frequently encoded by major genes (Oliva R & IL Quibod, 2017; Mundt CC, 2018). As described in the Introduction, the mechanisms of a few APR genes have been elucidated, such as Lr67, Lr34 and Yr36, which encode for a sugar transporter (Moore JW et al., 2015), an ATP-binding cassette transporter (Krattinger SG et al., 2009), and a detoxification protein (Fu D et al., 2009), respectively. Secondly, it could result from the fact that APR genes are rarely alone in a susceptible host genetic background but may be shielded by major genes. Finally, it could be due to the weaker selection pressure applied by APR genes on pathogens (since they allow some infection by wt pathogens by being only partially efficient and delayed in the season) (Mundt CC, 2018).

In the absence of relevant quantitative data concerning the first hypothesis, our parameterisation of the model gives the same mutation probability to overcome major genes and APR genes. Hence, the present study explores the latter two hypotheses. The possibility for APR genes to be shielded by major genes has been tested in Experiment 3 while the effect of selection pressure is highlighted by the difference between Experiments 1 and 2. The mutation probability to overcome the resistances was set at a high value, which could explain why, in our simulations, the combination of an APR gene with a major gene in a pyramided cultivar did not affect the durability of the APR gene in comparison to a cultivar that carried the APR gene only. Future work could investigate the potential of such pyramids with a lower mutation probability. On the other hand, our work emphasizes how shifts in selection pressure influence resistance durability. Indeed, APR genes were found to be very durable when they have a small efficiency and late activation. It may explain why some APR genes like Yr18, which has a small to moderate efficiency against stripe rust (Elahinia SA & JP Tewari, 2005; Qamar M et al., 2012) have shown high durability in the field (Krattinger SG et al., 2009). The efficiency of other APR genes like Lr12, Lr13, Lr22, Lr34, Lr35 and Lr37 have been measured between 80% and 90% against leaf rust (page 56 in Burdon JJ, 1987; McIntosh RA et al., 1995; Smale M et al., 1998). With such high efficiency, our simulations predicts that these genes could be quickly overcome. Nevertheless, depending on the age of resistance activation and the target pathogenicity trait, even if these genes were broken down, the resulting harsh competition between the different pathogen genotypes has the potential to provide some disease limitation, especially when deployed together with major resistance genes in mixture or rotation strategies. However, this conclusion strongly depends on the presence of fitness costs of pathogen adaptation to resistance (which influences the relative fitness of the wt and rb genotypes) as well as, likely, the dispersal abilities of the pathogen (which influences the migration of wt genotypes from susceptible to resistant hosts) and its mutation rate (which influences the appearance of rb genotypes). Furthermore, our results must be nuanced by the fact that we assumed that rb pathogens were penalised by a fitness cost on inactive APR genes, exactly as if the associated cultivars were susceptible. Experiments could be carried out in controlled conditions to test this hypothesis. We also assumed that APR genes switch suddenly from being inactive to active, whereas some rare available data rather indicate a gradual expression of APR genes (Ma H & RP Singh, 1996). Finally, while in our simulations, APR genes could target only one pathogenicity trait at a time, in the real world pathogenicity traits often vary in association (Parlevliet JE, 1979; Sache I & C de Vallavieille-Pope, 1995; Leclerc M et al., 2019). For example, Lr16-Lr18 targets latent period duration as well as sporulation rate and duration (Tomerlin JR et al., 1983) and Lr34-Yr18 affects both infection rate and latent period (Qamar M et al., 2012).

Regardless, our study represents a first attempt to numerically explore evolutionary and epidemiological outcomes of the deployment of adult plant resistance for the management of plant diseases. Furthermore, although this work was motivated by rust fungi of cereal crops, the generality of the model makes our results likely applicable to other pathosystems. Adult plant resistance has also been described in viruses (whilst rather called “mature plant resistance”). For instance, a cultivar of *Nicotiana edwardsonii*, activates a delayed monogenic resistance against *Tobacco mosaic virus, Tobacco necrosis virus* and *Tobacco bushy stunt virus* (Cole AB et al., 2004). Mature plant resistance has also been demonstrated in the greenhouse against *Cucumber mosaic virus* with a complete restriction of viral movement and systemic colonisation in mature bell pepper plants (Garcia-Ruiz H & JF Murphy, 2001) and against *Potato virus Y* with a restriction of tuber infection in potato (Kumar P et al., 2022).

## Supporting information

Raw data

## Acknowledgements

The authors thank Marta Zaffaroni and Jean-Loup Gaussen for their contribution to improve the R package *landsepi*, Loïc Houde for computing assistance and Jeremy Burdon for stimulating discussions. They also thank Timothée Poisot, Jean-Paul Soularue and an anonymous reviewer for their insightful comments on the manuscript.

## Data, script and code availability

The model is available in the open-access R package *landsepi* (Rimbaud L et al., 2018b); webpage: https://csiro-inra.pages.biosp.inrae.fr/landsepi/). Simulations were performed on the BioSP computational cluster from INRAE (https://biosp-cluster.mathnum.inrae.fr/). Simulation results are available in supplementary information.

## Supplementary information

**Text S1**. Full description of the mathematical model used in this study.

**Figure S1**. Heatmaps of the optimal pathogenicity trait targeted by an APR gene.

**Figure S2**. Heatmaps of the levels of evolutionary and epidemiological control, and average genotype frequencies in Experiment 2 when the target pathogenicity trait is the latent period duration.

**Figure S3**. Heatmaps of the levels of evolutionary and epidemiological control, and average genotype frequencies in Experiment 2 when the target pathogenicity trait is the sporulation rate.

**Figure S4**. Heatmaps of the levels of evolutionary and epidemiological control, and average genotype frequencies in Experiment 2 when the target pathogenicity trait is the sporulation duration.

**Figure S5**. Epidemiological outcome and dynamics of pathogen genotype frequencies in three examples of simulations.

**Figure S6**. Example of simulated fragmented landscapes used in Experiment 3.

**Figure S7**. Heatmaps of the levels of evolutionary and epidemiological control in Experiment 3 when the target pathogenicity trait is the latent period duration.

**Figure S8**. Heatmaps of the levels of evolutionary and epidemiological control in Experiment 3 when the target pathogenicity trait is the sporulation rate.

**Figure S9**. Heatmaps of the levels of evolutionary and epidemiological control in Experiment 3 when the target pathogenicity trait is the sporulation duration.

**Table S1**. Observed ranges of infection rate, latent period duration, sporulation rate and sporulation duration for rust fungi.

**Raw data**. Dataset of simulation results used in this study.

## Text S1

### Full description of the mathematical model used in this study

In the present work, the model is an adapted version of the one presented in Rimbaud L et al. (2018c), which simulates the clonal reproduction, spread and evolution of a pathogen in an agricultural landscape and over multiple cropping seasons. The original model simulated the deployment of resistant cultivars carrying major resistance genes or quantitative trait loci of resistance. In the present work, we introduced resistance genes with a delayed activation, i.e. Adult Plant Resistance (APR) genes. Cultivars that carry an APR gene are susceptible at the beginning of the cropping season and become resistant once the gene activates.

The model is described in the following, noting that we only reported equations that are relevant for the present study, i.e. equations related to qualitative resistance for which the pathogen and its host have a gene-for-gene interaction. Readers are referred to Rimbaud L et al. (2018c) for equations related to quantitative interactions. The model and its code are available in the R package *landsepi* (Rimbaud L et al., 2018b).

### Overview of the model

The model is stochastic, spatially-explicit and demo-genetic. It simulates the spread (through clonal reproduction and dispersal) and evolution (via mutation, selection and drift) of a plant pathogen in a heterogeneous landscape (the basic spatial unit is an individual field) across cropping seasons split by host harvests (which impose potential bottlenecks to the pathogen population). It is based on a spatial geometry for describing the landscape and allocation of different cultivars, a dispersal kernel for the dissemination of the pathogen, and a SEIR (‘Susceptible-Exposed-Infectious-Removed’ renamed HLIR for ‘healthy-latent-infectious-removed’ to avoid confusions with hosts that are genetically ‘susceptible’) structure with a discrete time step. The model simulates a wide array of deployment strategies: mosaics, mixtures, rotations and pyramiding of multiple resistance genes (i.e., in this work, major resistance genes or APR genes). These genes affect up to four pathogenicity traits: the infection rate, the duration of the latent period and the infectious period, and the propagule production rate. Resistance may be complete (i.e. complete inhibition of the targeted pathogenicity trait) or partial (i.e. the targeted pathogenicity trait is only softened), and activated from the beginning of the season, or later (APR). To each resistance gene in the host is associated a pathogenicity gene in the pathogen. Initially, the pathogen is not adapted to any source of resistance, and is only present on susceptible hosts. However, through mutation of pathogenicity genes, it can evolve and overcome resistance, possibly with penalty (fitness cost) on susceptible hosts. This model provides a useful tool to assess the performance of a wide range of deployment options via epidemiological and evolutionary outputs.

### Model assumptions

1. The spatial unit is a field, i.e. a piece of land delimited by boundaries and cultivated with a crop. The field is considered a homogeneous mixture of host individuals (i.e. there is no intra-field structuration).
2. Host individuals are in one of these four categories: H (healthy), L (latent, i.e. infected but not infectious nor symptomatic), I (infectious and symptomatic), or R (removed, i.e. epidemiologically inactive).
3. A host ‘individual’ is an infection unit and may correspond to a given amount of plant tissue (where a local infection may develop, e.g. fungal lesion) or a whole plant (e.g. systemic viral infection). In the first case, plant growth increases the amount of available plant tissue (hence the number of individuals) during the cropping season. Plant growth is deterministic (logistic growth) and only healthy hosts (state H) contribute to plant growth (castrating pathogen).
4. The decreasing availability of healthy host tissues (as epidemics spread) makes pathogen infection less likely (i.e. density-dependence due to plant architecture).
5. Host are cultivated (i.e. sown/planted and harvested), thus there is no host reproduction, dispersal or natural death.
6. Environmental and climate conditions are constant.
7. Components of a mixture are independent from each other (i.e. there is neither plant-plant interaction nor competition for space).
8. The pathogen is haploid and its reproduction mode is clonal.
9. Initially, the pathogen is not adapted to any source of resistance, and is only present on susceptible hosts (at state I).
10. Pathogen dispersal is isotropic (i.e. equally probable in every direction).
11. At the end of each cropping season, pathogens experience a bottleneck representing the off-season and then propagules are produced. Propagules are released at the first day of the following season.
12. Pathogenicity genes mutate independently from each other.
13. Pathogen adaptation to a given resistance gene consists in restoring the same pathogenicity trait as the one targeted by the resistance gene.
14. If a fitness cost penalizes pathogen adaptation to a given resistance gene, this cost is paid on hosts that do not carry this gene, and consists in a reduction in the same pathogenicity trait as the one targeted by the resistance gene. Fitness costs are multiplicative in case the pathogen carries multiple pathogenicity genes.
15. When there is a delay for activation of a given resistance gene (APR), the age of activation is the same for all hosts carrying this gene and located in the same field.
16. Variances of the durations of the latent and the infectious periods of the pathogen are not affected by plant resistance.

### Host-pathogen genetic interaction

The time steps of the model are indexed by t (t=1,…,TxY) where T is the number of time steps in a cropping season and Y the number of simulated years (i.e. cropping seasons).

#### Host genotype

A host genotype (indexed by v) is represented by a set of binary variables indicating the resistances genes it carries denoted as rg_g_ (g=1,…,G). The target pathogenicity trait of gene g is noted w(g)=e,γ,r,ϒ for infection rate, latent period duration, propagule production rate and infectious period duration, respectively. If gene rg_g_ is an APR gene, the age of resistance activation, t^a^, is drawn from a gamma distribution (a flexible continuous distribution from which durations in the interval [0; +∞[can be drawn) every year y (y=1,…,Y) and for every field i (i=1,…,J) planted with the cultivar carrying this gene. For convenience, this distribution is parameterised with the expectation, 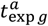, and variance, 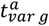, of the age of activation; and these are supposed equal (i.e. larger ages are also more variable):

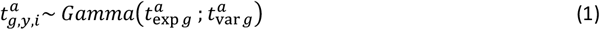

Note, the usual shape and scale parameters of a Gamma distribution, β_1_ and β_2_, can be calculated from the expectation and variance, exp and var, with: 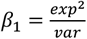 and 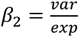, respectively.

Thus, for cultivar v carrying an APR gene g, this cultivar is susceptible between the beginning of a cropping season and t^a^-1 (i.e. rg_g_(v,t)=0), and resistant from t^a^ to the end of the cropping season (i.e. rg_g_(v,t)=1).

### Pathogen genotype

A pathogen genotype (indexed by p) is represented by a set of binary variables indicating the pathogenicity genes it carries, denoted as pg_g_.

### Interaction matrix

The interaction between host resistance genes and corresponding pathogenicity genes is represented by the following interaction matrix (denoted as INT^g^). This matrix shows the coefficients by which the value of the target pathogenicity trait (see further) is multiplied (except for latent period duration which varies in a direction opposite to that of the other traits: 1-ρ is replaced by 1+ρ and 1-θ is replaced by 1+θ):

| Infectivity matrix for gene $g$<br>$INT^g$ | | Host genotype $v$ | |
| --- | --- | --- | --- |
| | | Susceptible (SC)<br>$rg_g(v,t)=0$ | Resistant (RC)<br>$rg_g(v,t)=1$ |
| Pathogen<br>genotype $p$ | Non-adapted<br>$pg_g(p)=0$ | 1 | $1 - \rho_g$ |
| | adapted<br>$pg_g(p)=1$ | $1 - \theta_g$ | 1 |

with ρ_g_ the efficiency of resistance gene g on a non-adapted pathogen (ρ_g_ = 1 for a typical major resistance gene conferring immunity, but ρ_g_ < 1 for partial resistance), and θ_g_ the fitness cost paid by an adapted pathogen on a host that does not carry the resistance gene (θ_g_=0 corresponds to absence of cost, θ_g_=1 corresponds to loss of pathogenicity on the concerned host).

### Host and pathogen demo-genetic dynamics

In the following, H_i,v,t_, L_i,v,p,t_, I_i,v,p,t_, R_i,v,p,t_, and Pr_i,p,t_ respectively denote the number of healthy, latent, infectious and removed host individuals, and pathogen propagules, respectively, in field i (i=1,…,J), for cultivar v (v=1,…,V), pathogen genotype p (p=1,…,P) at time step t (t=1,…,TxY).

#### Host growth

Only healthy hosts (denoted as *H*_*i,v,t*_) are assumed to contribute to the growth of the crop. Thus, at each step t during a cropping season, the plant cover of cultivar v in field i increases as a logistic function, and the new amount of healthy plant tissue is:

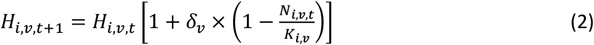

with δ_v_ the growth rate of cultivar 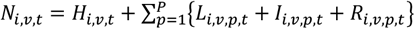 the total number of hosts in field i for cultivar v and at time t and 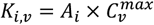 the carrying capacity of cultivar v in field i, which depends on A_i_, the area of the field, and 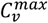, the maximal density for cultivar v. Note that when a mixture of several cultivars is present in a given field, decreased growth due to susceptible plants being diseased is not compensated for by increased growth of resistant plants.

#### Contamination of healthy hosts

The healthy compartment (H) is composed of hosts that are free of pathogen propagules (H^1^), as well as hosts contaminated (but not yet infected) by the arrival of such propagules (H^2^). At time t in field i and for cultivar v, the number of contaminable hosts (i.e. accessible to pathogen propagules, denoted as 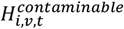) depends on the proportion of healthy hosts 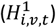 in the host population (*N*_*i,v,t*_):

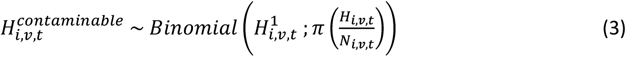

with 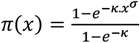, a sigmoid function with π(0)=0 and π(1)=1, giving the probability for a healthy host to be contaminated. Here, we assume that healthy hosts are not equally likely to be contacted by propagules, for instance because of plant architecture. Moreover, as the local severity of disease increases, eventually the probability for a single propagule to contaminate a healthy host declines due to the decreased availability of host tissues.

Following the arrival of propagules of pathogen genotype p in field i at time t (denoted as 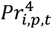, see below), susceptible hosts become contaminated. The pathogen genotypes of these propagules are distributed among contaminable hosts according to their proportional representation in the total pool of propagules. Thus, for cultivar v, the vector describing the maximum number of contaminated hosts by each pathogen genotype (denoted as 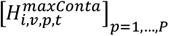) is given by a multinomial draw:

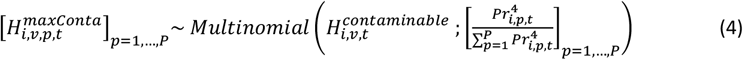

However, the number of deposited propagules 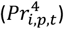 may be smaller than the maximal number of contaminated hosts 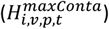. Thus, the true number of hosts of cultivar v, contaminated by pathogen genotype p in field i at t (denoted as 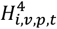) is:

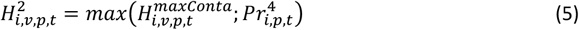

#### Infection

Between t and t+1, in field i, contaminated hosts 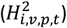 become infected (state L) with probability e_v,p_, which depends on the maximum expected infection efficiency, e_max_, and the interaction between host (v) and pathogen (p) genotypes:

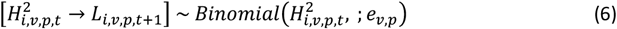

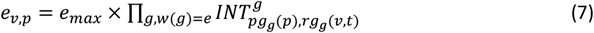

#### Latent period

Infected hosts become infectious (state I) after a latent period (LI) drawn from a Gamma distribution parameterised with an expected value, γ_exp v,p_, and variance, γ_var_, similar to the age of resistance activation. The expected duration of the latent period depends on the minimal expected duration, γ_min_, and the interaction between host (v) and pathogen (p) genotypes:

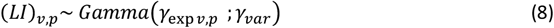

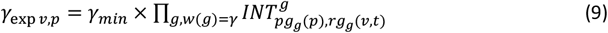

#### Infectious period

Finally, infectious hosts become epidemiologically inactive (i.e. they no longer produce propagules and thus are in state R, ‘removed’) after an infectious period (IR) drawn from a Gamma distribution parameterised with expected value, ϒ _exp v,p_ and variance, ϒ _var_, similar to the latent period. The expected duration of the infectious period depends on the maximal expected duration, ϒ _max_, and the interaction between host (v) and pathogen (p) genotypes:

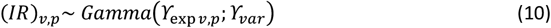

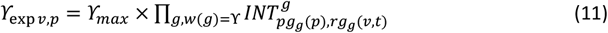

#### Pathogen reproduction

In field i at time t, infectious hosts associated with pathogen genotype p produce a total number of propagules (denoted as 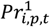), drawn from a Poisson distribution whose expectation, r_exp v,p_, depends on the maximal expected number of propagules produced by a single infectious host per time step, r_max_, and the interaction between host (v) and pathogen (p) genotypes:

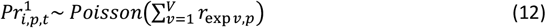

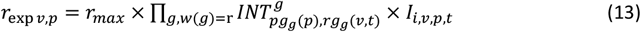

#### Pathogen mutation

The following algorithm is repeated independently for every pathogenicity gene g:

1. the pathotype (i.e. the level of adaptation with regard to gene g, indexed by q; q=1,…,Q_g_; with Q_g_=2 since the pathotype is either adapted, or non-adapted) of the pathogen propagules is retrieved from their genotype p;
2. propagules can mutate from pathotype q to pathotype q’ with probability 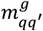 such as 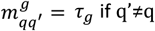 if q’≠q (hence 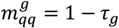 since Q_g_=2). Thus, in field i at time t, the vector of the number of propagules of each pathotype arising from pathotype q (denoted as 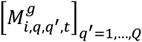) is given by a multinomial draw:

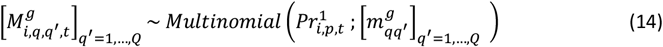
3. the total number of propagules belonging to pathotype q’ and produced in field i at time t (denoted as 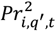) is:

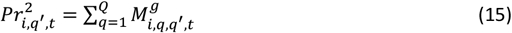
4. the new propagule genotype p’ is retrieved from its new pathotype (q’), and the number of propagules is incremented using a variable denoted as 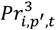.

In this model, it should be noted that the mutation probability τ_g_ is not the classic ‘mutation rate’ (i.e. the number of genetic mutations per generation per base pair), but the probability for a propagule to change its pathogenicity on a resistant cultivar carrying gene g. This probability depends on the classic mutation rate, the number and nature of the specific genetic mutations required to overcome gene g, and the potential dependency between these mutations.

#### Pathogen dispersal

Propagules can migrate from field i (whose area is A_i_) to field i’ (whose area is A_i’_) with probability μ_ii’_, computed from:

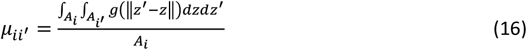

with ‖*z*^′^ − *z*‖ the Euclidian distance between locations z and z’ in fields i and i’, respectively, and g(.) the two-dimensional dispersal kernel of the propagules. Thus, at time t, the vector of the number of propagules of genotype p migrating from field i to each field i’ (denoted as 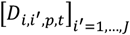) is:

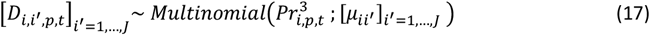

and the total number of propagules arriving in field i’ at time t (denoted as 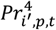) is:

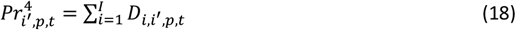

We consider that propagules landing outside the boundaries of the simulated landscape are lost (absorbing boundary condition), and there are no propagule sources external to the simulated landscape.

#### Seasonality

Let *t*^0^(*y*) and *t*^*f*^(*y*) denote the first and last days of cropping season y (y=1,…,Y), respectively. The plant cover in field i for cultivar v at the beginning of cropping season y is set at 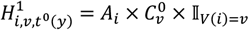, with 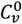 the plantation density of cultivar v and 𝕀_*V* (*i*)_ an indicative variable set at 1 when field i is cultivated with cultivar v and 0 otherwise. At the end of a cropping season, the host is harvested. We assume that the pathogen needs a green bridge to survive the off-season. This green bridge could, for example, be a wild reservoir or volunteer plants remaining in the field. The size of this reservoir imposes a bottleneck for the pathogen population. The number of remaining infected hosts in field i for cultivar v and pathogen genotype p (denoted by 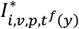) at the end of the off-season is given by:

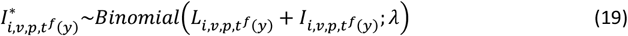

with λ the survival probability of infected hosts. Depending on host (v) and pathogen (p) genotypes, the number of propagules produced by the remaining hosts during their whole infectious period (denoted by 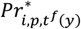) is drawn from a Poisson distribution:

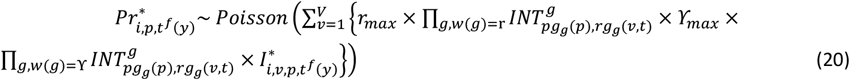

These propagules can mutate and disperse exactly as happens during the cropping season and constitute the initial inoculum for the next cropping season.

#### Initial conditions

At the beginning of a simulation, healthy hosts are planted in each field as previously described. The initial pathogen population is assumed to be fully non-adapted to the resistance host cultivars, and is only present in susceptible fields with probability ϕ. Then the initial number of infectious hosts in these fields is: 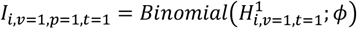.

### Model outputs

The model utilises multiple criteria to enable the assessment of different deployment strategies with regard to epidemiological, evolutionary and economic outcomes. Here we detail only outputs relevant to the current study.

#### Epidemiological output

The ability of a given deployment strategy to reduce disease impact on the resistant cultivar(s) is measured by the relative green leaf area (GLA), i.e., the proportion of healthy hosts relative to the total number of hosts, averaged for every cultivar v across the whole simulation run:

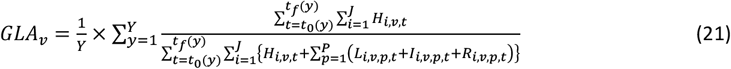

#### Evolutionary outputs

Resistance durability measures the ability of a given deployment strategy to limit pathogen evolution and delay resistance breakdown. Durability is evaluated using the time t* when the number of resistant hosts infected by adapted genotypes (e.g., if v=2 for the resistant cultivar and p=2 for the rb pathogen, 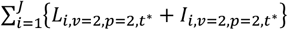) exceeds a threshold above which extinction of this strain is unlikely (fixed at 50,000, see Rimbaud L et al., 2018c, supporting Text S2 for details). To understand the contribution of the different pathogen genotypes to an epidemic, we also calculate, across the whole simulation run and for every cultivar, the proportion of infections due to each pathogen genotype relative to all infections. The contribution of pathogen p to epidemics on cultivar v is computed as follows:

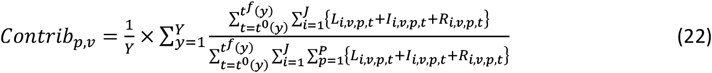

**Figure S1.**
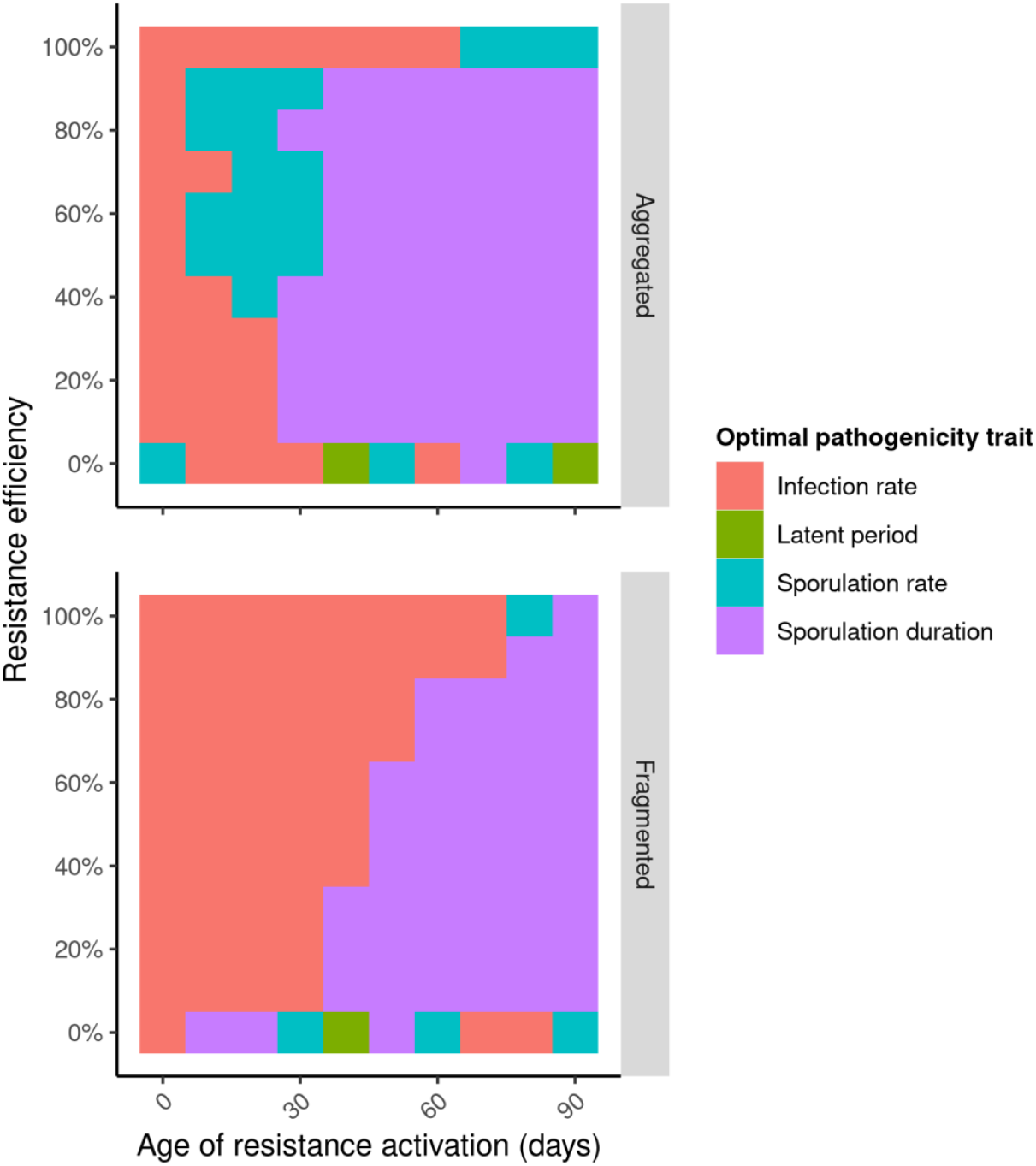
Heatmaps indicating the optimal pathogenicity trait targeted by an APR gene with respect to the level of epidemiological control (i.e., disease limitation, measured by the Green Leaf Area, ‘GLA’) on the resistant cultivar in the absence of pathogen evolution for different levels of resistance efficiency (vertical axis) and age of resistance activation (horizontal axis), for strong (top) or weak (bottom) levels of spatial aggregation.

**Figure S2.**
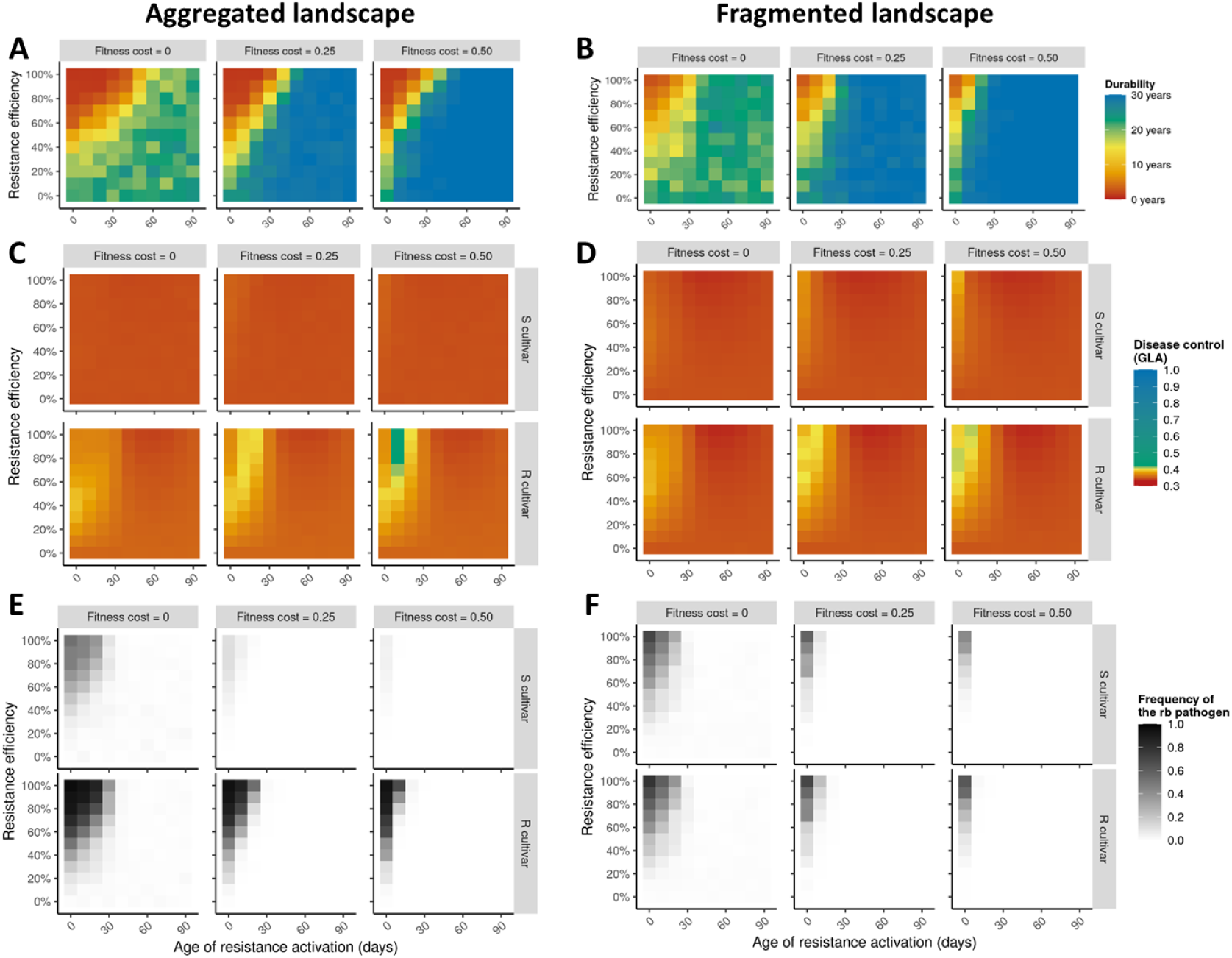
Heatmaps of the levels of evolutionary control (resistance durability as measured by the number of years before the emergence of the resistance-breaking (‘rb’) pathogen genotype, panels A and B), epidemiological control (i.e. disease limitation, measured by the Green Leaf Area (‘GLA’) on the susceptible (‘S’) and the resistant (‘R’) cultivars, panels C and D) and average frequency of the rb pathogen (panels E and F) for different levels of resistance efficiency (vertical axis), age of resistance activation (horizontal axis) and fitness cost of pathogen adaptation (columns), for strong (panels A, C, E) or weak (B, D, F) levels of spatial aggregation. The target pathogenicity trait is the latent period duration.

**Figure S3.**
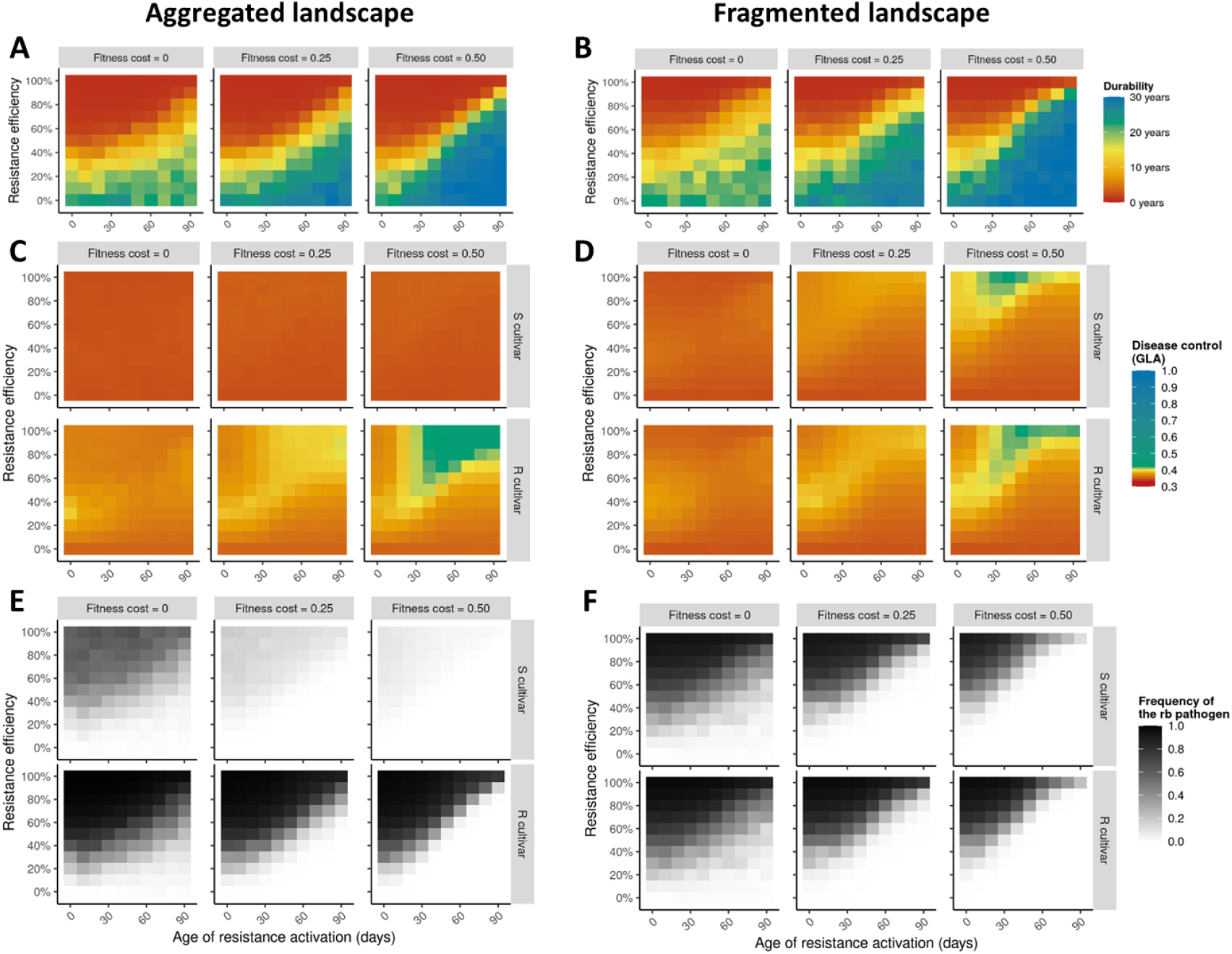
Heatmaps of the levels of evolutionary control (resistance durability as measured by the number of years before the emergence of the resistance-breaking (‘rb’) pathogen genotype, panels A and B), epidemiological control (i.e. disease limitation, measured by the Green Leaf Area (‘GLA’) on the susceptible (‘S’) and the resistant (‘R’) cultivars, panels C and D) and average frequency of the rb pathogen (panels E and F) for different levels of resistance efficiency (vertical axis), age of resistance activation (horizontal axis) and fitness cost of pathogen adaptation (columns), for strong (panels A, C, E) or weak (B, D, F) levels of spatial aggregation. The target pathogenicity trait is the sporulation rate.

**Figure S4.**
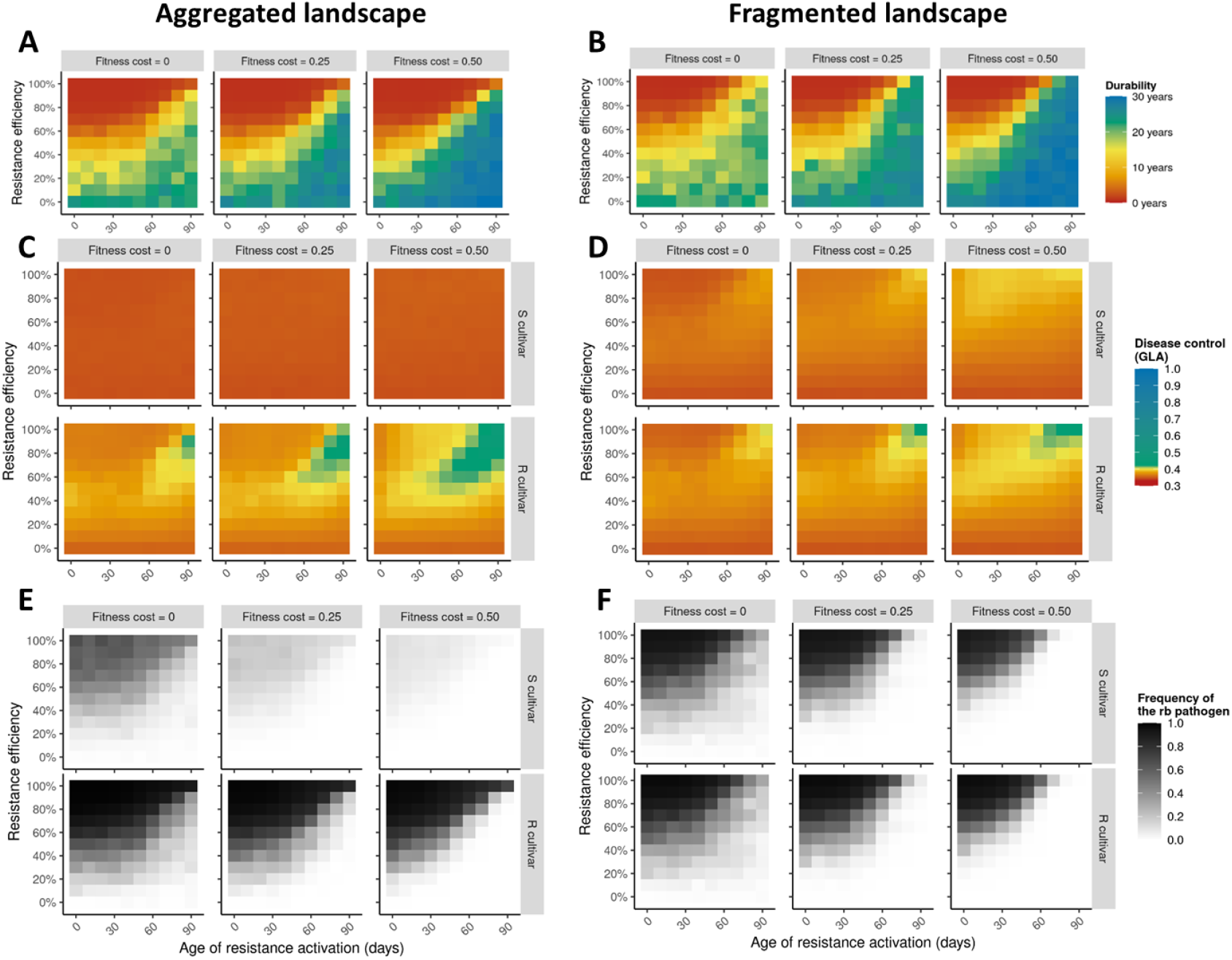
Heatmaps of the levels of evolutionary control (resistance durability as measured by the number of years before the emergence of the resistance-breaking (‘rb’) pathogen genotype, panels A and B), epidemiological control (i.e. disease limitation, measured by the Green Leaf Area (‘GLA’) on the susceptible (‘S’) and the resistant (‘R’) cultivars, panels C and D) and average frequency of the rb pathogen (panels E and F) for different levels of resistance efficiency (vertical axis), age of resistance activation (horizontal axis) and fitness cost of pathogen adaptation (columns), for strong (panels A, C, E) or weak (B, D, F) levels of spatial aggregation. The target pathogenicity trait is the sporulation duration.

**Figure S5.**
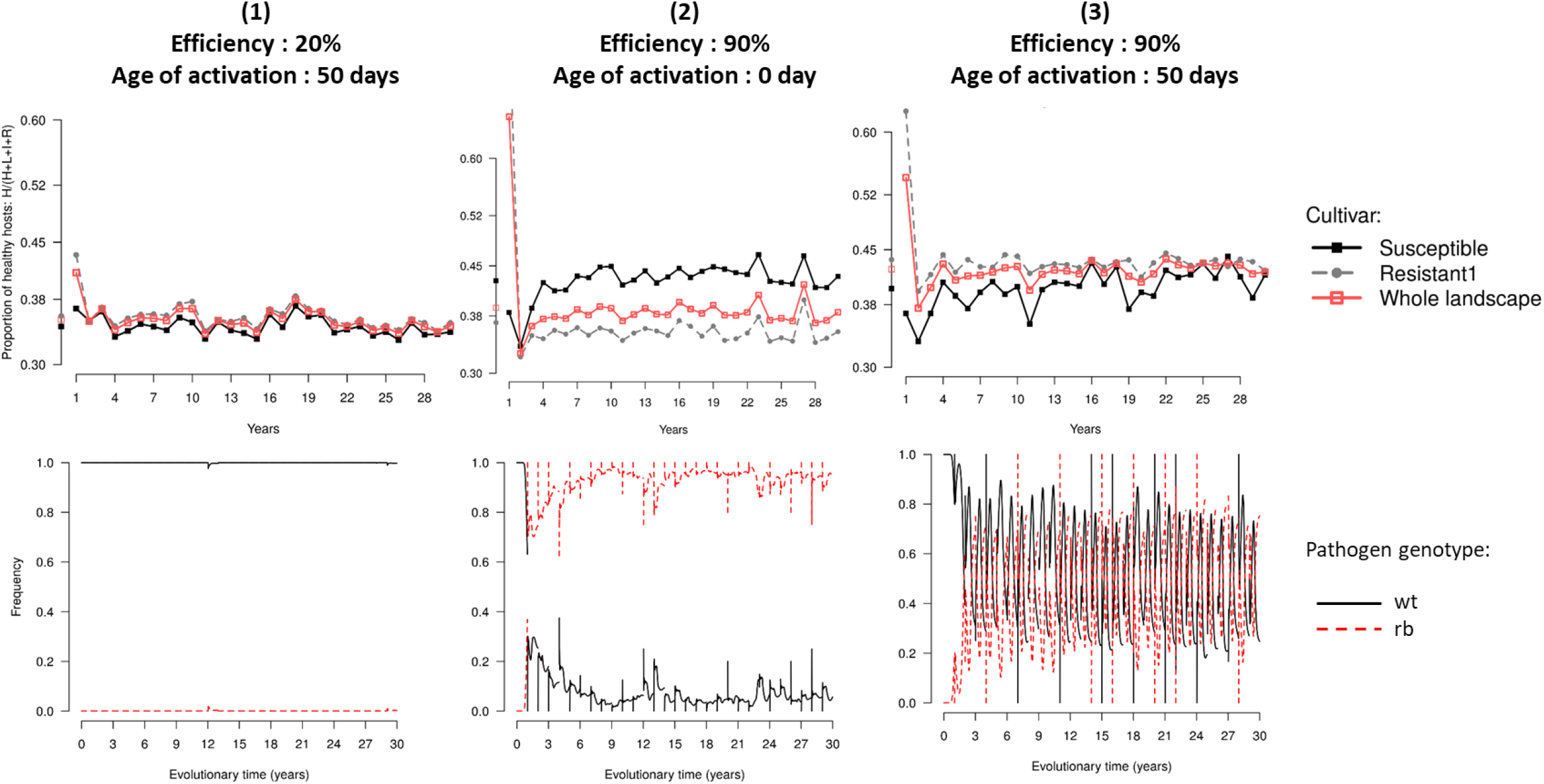
Epidemiological outcome (represented by the relative Green Leaf Area, top line) and dynamics of pathogen genotype frequencies (bottom line, ‘wt’ refers to the wild-type and ‘rb’ to the resistance-breaking pathogen genotype) in three examples of simulations where a single APR is deployed in a susceptible landscape with low level of spatial aggregation. Situations 1, 2 and 3 are pointed in Figure 3. The pathogenicity trait targeted by resistance is the infection rate and the fitness cost of adaptation is θ=0.50.

**Figure S6.**
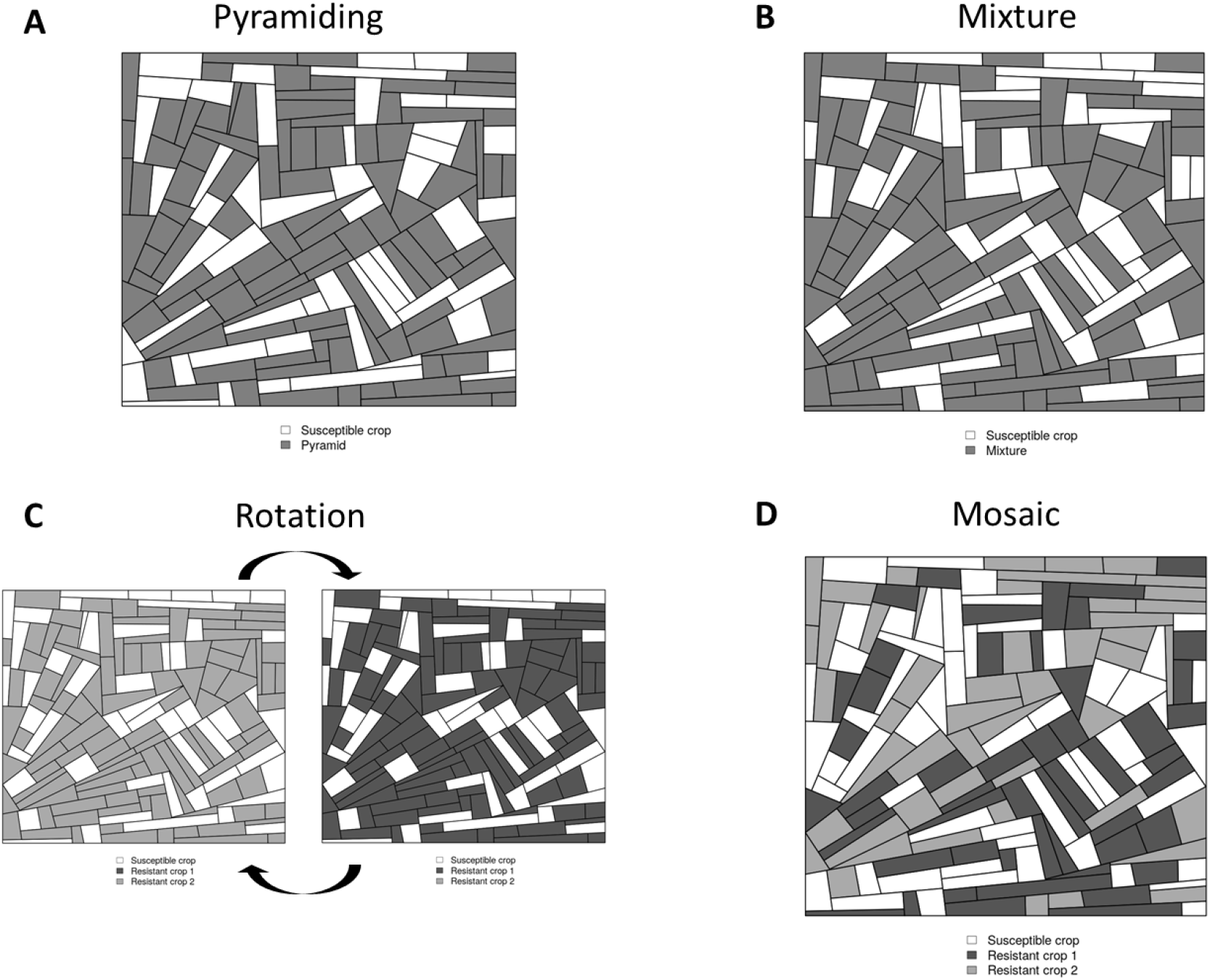
Example of simulated fragmented landscapes used in Experiment 3 (APR + MG). For all deployment strategies, 1/3 of the landscape was composed of the susceptible cultivar. The remaining 2/3 were occupied either by: A) a single cultivar carrying the two genes (pyramid strategy); B) a mixture (in every field) of two resistant cultivars in balanced proportions (each cultivar carrying one of the two genes); C) a rotation of these two resistant cultivars (every year); or D) a mosaic of the two resistant cultivars in balanced proportions (every cultivar representing 1/3 of the landscape area).

**Figure S7.**
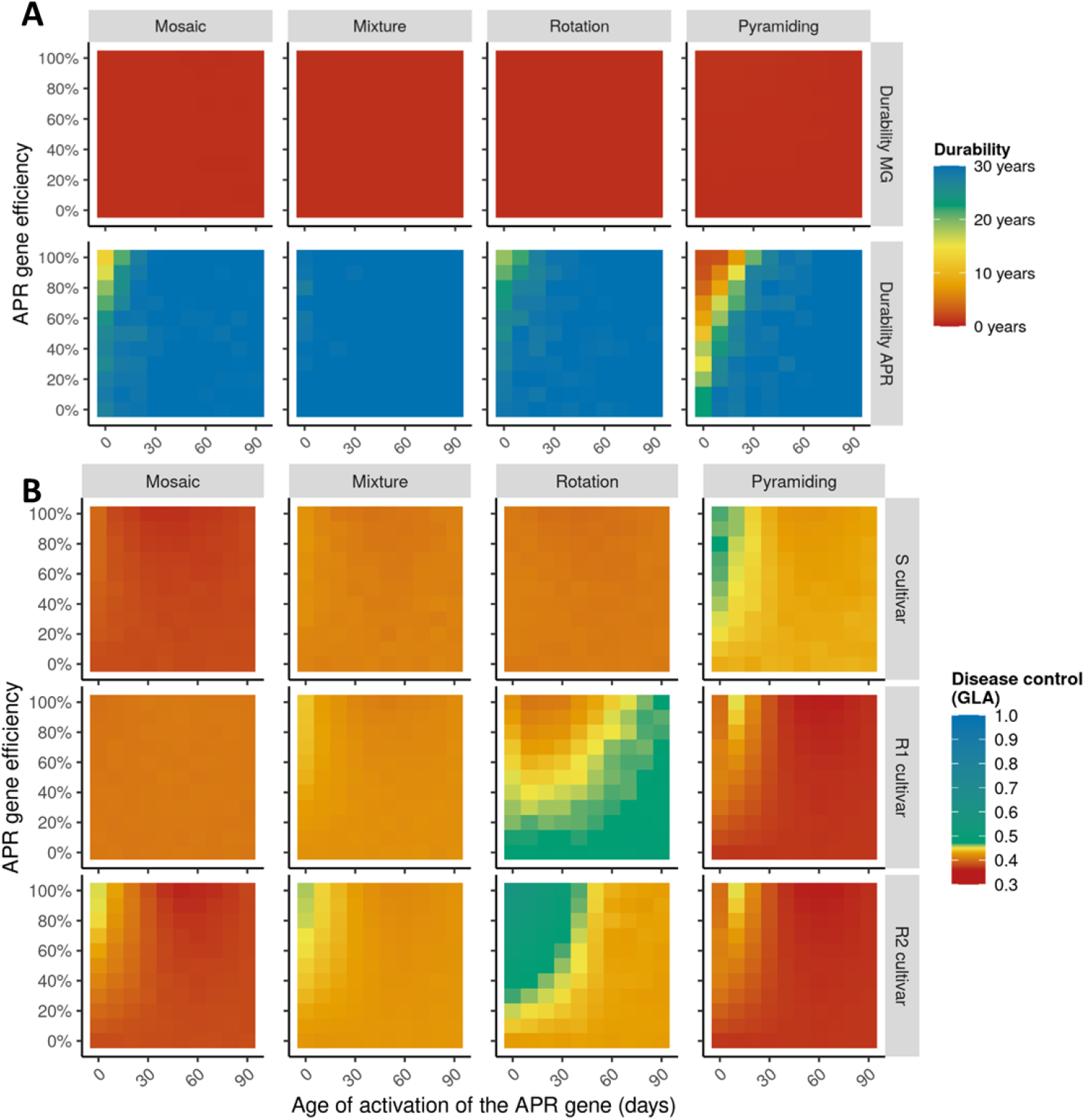
Heatmaps showing the levels of A) evolutionary control (resistance durability, measured by the number of years before the emergence of resistance-breaking genotypes) and B) epidemiological control (i.e., disease limitation, measured by the Green Leaf Area, ‘GLA’) on a susceptible cultivar ‘S’, a resistant cultivar ‘R1’ carrying a completely efficient major gene (‘MG’) and a resistant cultivar ‘R2’ carrying an APR gene, for different levels of APR efficiency (vertical axis), age of APR activation (horizontal axis) and deployment strategies (columns; note that for pyramiding, R1 and R2 refer to the same cultivar). The target pathogenicity trait of the APR gene is the latent period duration, the level of spatial aggregation is low, and the fitness cost is 0.50.

**Figure S8.**
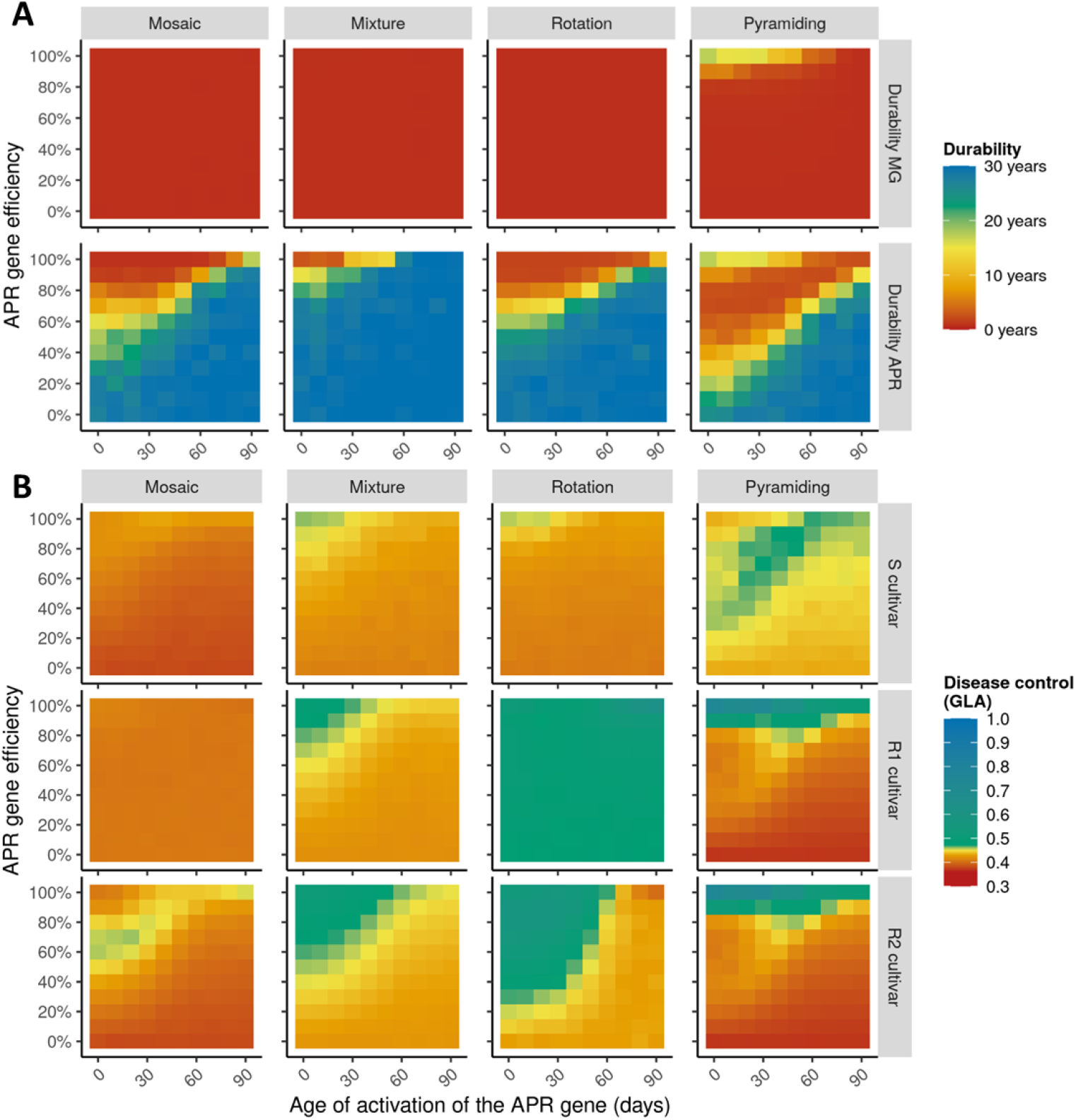
Heatmaps showing the levels of A) evolutionary control (resistance durability, measured by the number of years before the emergence of resistance-breaking genotypes) and B) epidemiological control (i.e., disease limitation, measured by the Green Leaf Area, ‘GLA’) on a susceptible cultivar ‘S’, a resistant cultivar ‘R1’ carrying a completely efficient major gene (‘MG’) and a resistant cultivar ‘R2’ carrying an APR gene, for different levels of APR efficiency (vertical axis), age of APR activation (horizontal axis) and deployment strategies (columns; note that for pyramiding, R1 and R2 refer to the same cultivar). The target pathogenicity trait of the APR gene is the sporulation rate, the level of spatial aggregation is low, and the fitness cost is 0.50.

**Figure S9.**
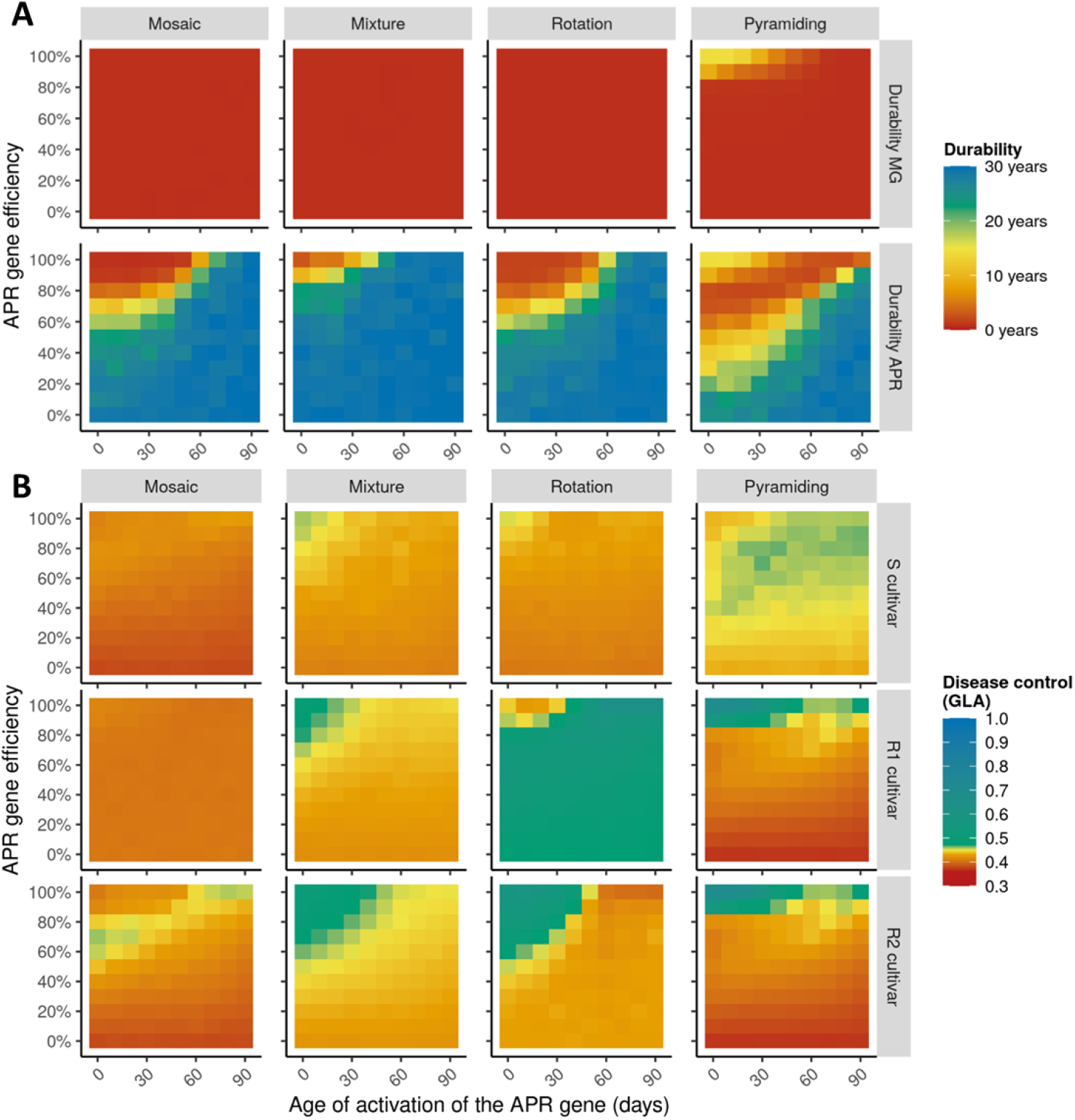
Heatmaps showing the levels of A) evolutionary control (resistance durability, measured by the number of years before the emergence of resistance-breaking genotypes) and B) epidemiological control (i.e., disease limitation, measured by the Green Leaf Area, ‘GLA’) on a susceptible cultivar ‘S’, a resistant cultivar ‘R1’ carrying a completely efficient major gene (‘MG’) and a resistant cultivar ‘R2’ carrying an APR gene, for different levels of APR efficiency (vertical axis), age of APR activation (horizontal axis) and deployment strategies (columns; note that for pyramiding, R1 and R2 refer to the same cultivar). The target pathogenicity trait of the APR gene is the sporulation duration, the level of spatial aggregation is low, and the fitness cost is 0.50.

**Table S1.**
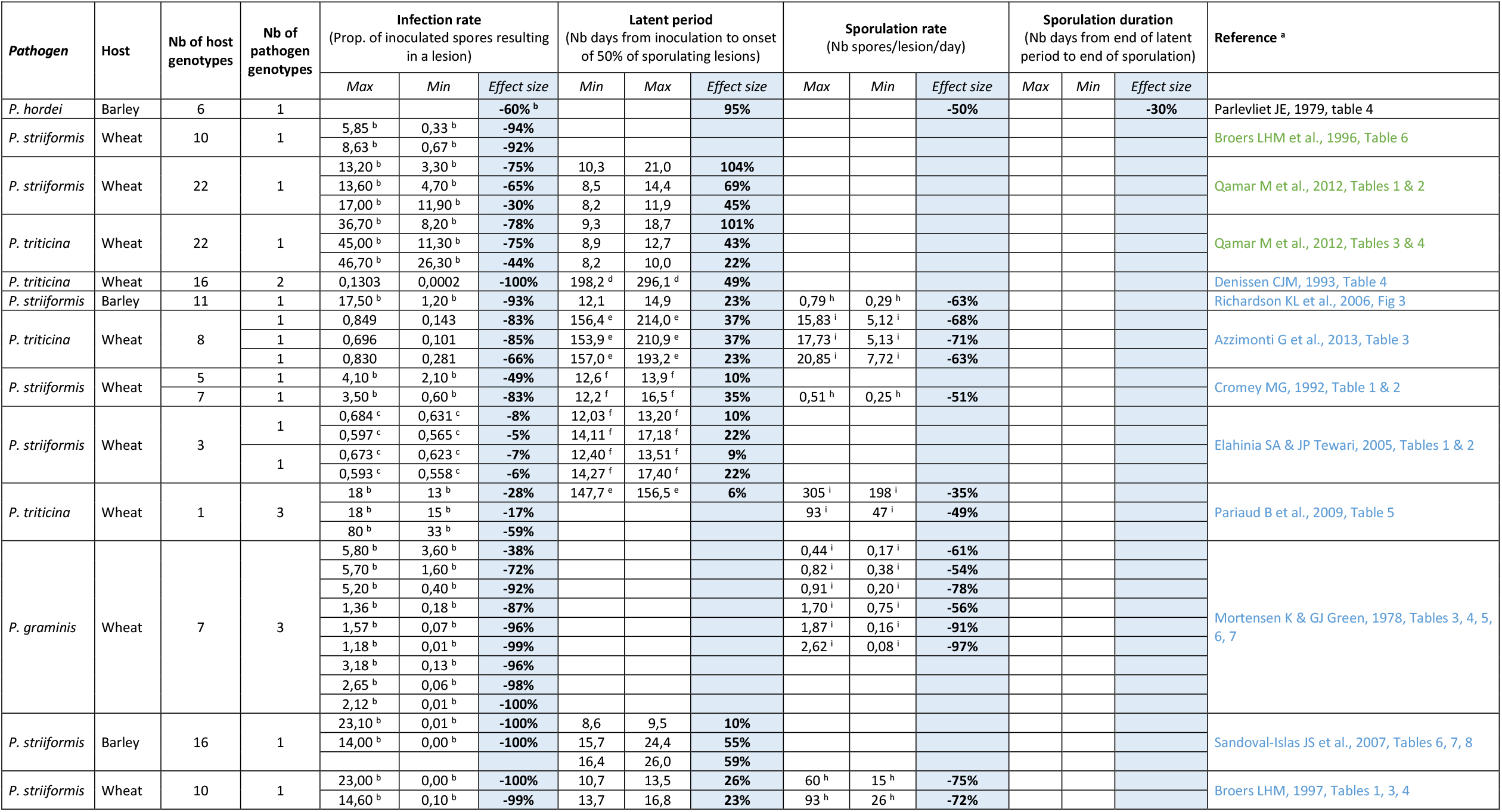
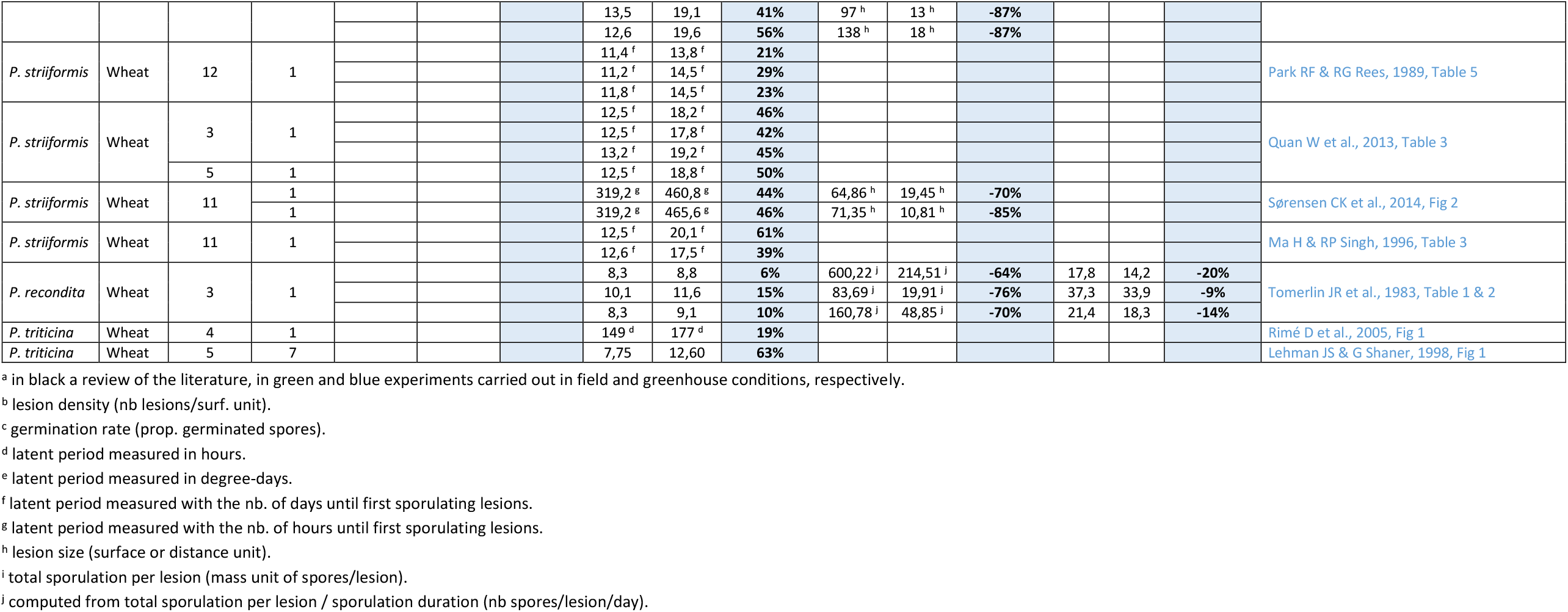
Observed ranges of infection rate, latent period duration, sporulation rate and sporulation duration for rust fungi (genus *Puccinia*) measured in different cultivars of wheat and barley. For a given study, different lines refer to different trials carried out in different conditions (time, temperature, pathogen genotype, leaf stage). Footnotes indicate when the measured variable is not exactly the same as the one used in the *landsepi* model (and of which a definition is given in the first line of the table).

