## Supplementary material for "Durable resistance or efficient disease control? Adult Plant Resistance (APR) at the heart of the dilemma": Raw data: readme.txt

########################    RAW DATA   #####################

***     Durable resistance or efficient disease control?     ***
*** Adult Plant Resistance (APR) at the heart of the dilemma ***

by: Loup Rimbaud, Julien Papaïx, Jean-François Rey, Benoît Moury, Luke Barrett, Peter Thrall

All simulations are computed using the R package landsepi
Package webpage: https://csiro-inra.pages.biosp.inrae.fr/landsepi/
To download the package: https://cran.r-project.org/web/packages/landsepi/index.html

Raw data used for this study are stored in three folders corresponding to
the three numerical experiments (APR_1, APR_2 and APR_3, respectively). 

Every folder includes: 
- an R script for the generation of a simulation plan
- 10 folders containing simulation results for 10 stochastic replicates
- an R script for the analysis of the results and generation of graphics

Every folder containing the simulation results includes:
- a text file "myDesign.txt" containing the simulation plan and outputs (1 line per simulation)
- a text file "parameters.txt" containing a summary of all parameter values
- a text file "param_disp_patho.txt" containing the dispersal matrix of the pathogen
- a text file "param_landscape.txt" containing the landscape structure 
  (i.e. for every polygon [lines] and every year [columns] the index of the cultivated host)

################################################################
