## Supplementary material for "Durable resistance or efficient disease control? Adult Plant Resistance (APR) at the heart of the dilemma": Raw data: simul_plan_APR_1.R

```
library("landsepi", lib.loc="/home/lrimbaud/R/library")

## NUMERICAL DESIGN
myDesign <- expand.grid(id_landscape = 1:5
                        , aggreg = c(0.05, 3)
                        , fitness_cost = c(0.5)
                        , mutation_prob = c(0)
                        , R_efficiency = seq(0,1,0.1)
                        , time_to_activ_exp = seq(0,90,10)
                        , target_trait = c("IR", "LAT", "PR", "IP"))

n <- nrow(myDesign)
myDesign <- cbind(simul = 1:n, seed = sample(n*100, n), myDesign)


## Simulation parameters
simul_params <- createSimulParams()
simul_params <- setTime(simul_params, Nyears = 30, nTSpY = 120)

## Pathogen parameters
basic_patho_param <- loadPathogen("rust")
simul_params <- setPathogen(simul_params, basic_patho_param)     
## Initial conditions 
simul_params <- setInoculum(simul_params, 5E-4)

gene1 <- loadGene(name = "gene1", type = "majorGene")
genes <- data.frame(gene1, stringsAsFactors = FALSE)

## SIMULATIONS
for (i in 1:n){  
    print(paste("Running simulation", i, "/", n))
    
    ## seed
    simul_params <- setSeed(simul_params, myDesign$seed[i])
    ## landscape
    landscape <- loadLandscape(myDesign$id_landscape[i])
    simul_params <- setLandscape(simul_params, landscape)
    ## Dispersal
    disp_patho <- loadDispersalPathogen(myDesign$id_landscape[i])
    simul_params <- setDispersalPathogen(simul_params, disp_patho)
    
    ## Cultivars
    cultivar1 <- loadCultivar(name = "Susceptible", type = "growingHost")
    cultivar2 <- loadCultivar(name = "Resistant", type = "growingHost")
    cultivars <- data.frame(rbind(cultivar1, cultivar2), stringsAsFactors = FALSE)      
    
    simul_params <- setCultivars(simul_params, cultivars)  
    
    ## Allocate cultivars to croptypes
    croptypes <- loadCroptypes(simul_params, names = c("Susceptible crop", "Resistant crop"))
    croptypes <- allocateCroptypeCultivars(croptypes, "Susceptible crop", "Susceptible")
    croptypes <- allocateCroptypeCultivars(croptypes, "Resistant crop", "Resistant")
    
    simul_params <- setCroptypes(simul_params, croptypes)
    
    
    ## Genes
    genes$efficiency <- myDesign$R_efficiency[i]  ## update resistance efficiency
    genes$time_to_activ_exp <- myDesign$time_to_activ_exp[i]  ## update time to activation
    genes$time_to_activ_var <- myDesign$time_to_activ_exp[i]  ## update time to activation
    genes$mutation_prob <- myDesign$mutation_prob[i]  ## update mutation prob
    genes$fitness_cost <- myDesign$fitness_cost[i]  ## update fitness cost
    genes$target_trait <- as.character(myDesign$target_trait[i])  ## update target trait
    
    genes$Nlevels_aggressiveness <- 1
    if (as.character(myDesign$target_trait[i])=="LAT"){
        genes$efficiency <- genes$efficiency / 2
    }
    
    
    simul_params <- setGenes(simul_params, genes)
    
    ## Allocate genes to cultivars
    simul_params <- allocateCultivarGenes(simul_params, "Resistant", c("gene1"))
    
    ## Allocate croptypes to landscape  
    simul_params <- allocateLandscapeCroptypes(simul_params,
                                               rotation_period = 0,
                                               rotation_sequence = croptypes$croptypeID,
                                               rotation_realloc = FALSE,
                                               prop = c(1/3, 2/3),
                                               aggreg = myDesign$aggreg[i],
                                               graphic = FALSE)

    
    ## Outputs
    outputList <- loadOutputs(epid_outputs = c("audpc","audpc_rel","gla","gla_rel"), evol_outputs = "")
    simul_params <- setOutputs(simul_params, outputList)
    
    ## run simulation
    res <- runSimul(simul_params, writeTXT = FALSE, graphic = FALSE)

    ## Extract outputs
    contrib <- res$epid_outputs$contrib[,,"Whole simulation"]
    myDesign[i,c("audpc_S")] <- mean(res$epid_outputs$audpc$Susceptible, na.rm=TRUE)
    myDesign[i,c("audpc_rel_S")] <- mean(res$epid_outputs$audpc_rel$Susceptible, na.rm=TRUE)
    myDesign[i,c("gla_S")] <- mean(res$epid_outputs$gla$Susceptible, na.rm=TRUE)
    myDesign[i,c("gla_rel_S")] <- mean(res$epid_outputs$gla_rel$Susceptible, na.rm=TRUE)
    myDesign[i,c("audpc_R")] <- mean(res$epid_outputs$audpc$Resistant, na.rm=TRUE)
    myDesign[i,c("audpc_rel_R")] <- mean(res$epid_outputs$audpc_rel$Resistant, na.rm=TRUE)
    myDesign[i,c("gla_R")] <- mean(res$epid_outputs$gla$Resistant, na.rm=TRUE)
    myDesign[i,c("gla_rel_R")] <- mean(res$epid_outputs$gla_rel$Resistant, na.rm=TRUE)
    myDesign[i,c("audpc_tot")] <- mean(res$epid_outputs$audpc$total)
    myDesign[i,c("audpc_rel_tot")] <- mean(res$epid_outputs$audpc_rel$total, na.rm=TRUE)
    myDesign[i,c("gla_tot")] <- mean(res$epid_outputs$gla$total, na.rm=TRUE)
    myDesign[i,c("gla_rel_tot")] <- mean(res$epid_outputs$gla_rel$total, na.rm=TRUE)
    

    ## Create myDesign.txt at first simulation and then append
    if (i ==1){
        write.table(myDesign[i,], paste(simul_params@OutputDir, "/myDesign.txt", sep="")
                    , append=FALSE, row.names = FALSE, col.names = TRUE)      
    } else {
        write.table(myDesign[i,], paste(simul_params@OutputDir, "/myDesign.txt", sep="")
                    , append=TRUE, row.names = FALSE, col.names = FALSE)
    }
}
```
