## Supplementary material for "Durable resistance or efficient disease control? Adult Plant Resistance (APR) at the heart of the dilemma": Raw data: analyseAPR_2.R

```
######################################################
#########            analyseAPR_2.R             ########
######################################################
#### only one resistance gene
## landscape : Mosaic S + R[GFG], 1/3-2/3, Nyears = 30 years
## Variation of target trait (4 values), Reff & timeToR (11 values for 0 to 1)
## for different fitness cost (0, 0.25, 0.5), level of aggregation (0.05, 3)
## Stochastic replicates: 5 landscapes, 10 replicates per landscape
## Evolutionary outputs: dur_gene1.
## Epidemiological outputs: audpc, audpc_rel, gla, gla_rel, contrib_patho1, contrib_patho2 for S, R and tot
## No need for audpc and gla in whole landscape because:
## (audpc_tot = audpc_S + audpc_R     and    audpc_rel_tot almost = mean(audpc_rel_S, audpc_rel_R))
## Strong correlation between absolute and relative outputs 
## prefer GLA because higher variation
## --> use relative outputs for easier comparison between S and R (uneven proportions)
## contrib_patho1 = 1 - contrib_patho2 (except in case of pathogen absence)


rm(list=ls())
graphics.off()

run = "APR_2"


########  GRAPHICAL PACKAGES  ########
library("ggplot2")     ## for function ggplot
library("lemon")       ## for facet_rep_grid()
library("reshape2")     ## for melt
library("scales")       ## for rescale
library("PerformanceAnalytics")  ## for chart.Correlation
source("/home/loup/Dropbox/scriptsLPR/graphics_lpr.r") 
cb_palette <- c("#000000", "#999999", "#E69F00", "#56B4E9", "#009E73", "#F0E442", "#0072B2", "#D55E00", "#CC79A7", "#b71c1c")
names(cb_palette) <- c("black", "grey", "orange", "cyan", "green", "yellow", "blue", "red", "purple", "darkred")
######################################


######################
####     DATA     ####
######################
setwd(paste("/home/loup/Documents/model_resistance/RES/simul", run, sep=""))

## Load data
allFiles <- list.files(getwd())
filesToOpen <- allFiles[grep("simul_landsepi_2022-07-08", allFiles)]

mydata <- data.frame()
for (i in 1:length(filesToOpen)){
    data_tmp <- read.table(paste(filesToOpen[i], "/myDesign.txt", sep=""), sep=" ", dec=".",header=TRUE)
    data_tmp[,1] <- i
    colnames(data_tmp)[1] <- "rep"
    mydata <- rbind(mydata, data_tmp)
}

## Transform into factors
for (k in c("rep", "id_landscape","fitness_cost","mutation_prob"))
    mydata[,k] <- factor(mydata[,k])
traits <- c("IR", "LAT", "PR", "IP")
names(traits) <- c("Infection rate", "Latent period"
                   , "Sporulation rate", "Sporulation duration")
mydata$target_trait <- factor(mydata$target_trait, levels=traits)
mydata$aggreg <- factor(mydata$aggreg, levels=c(3, 0.05))

## Transform durability in % of the maximum possible value (i.e. 30 years) 
nYears <- 30
nTSpY <- 120
mydata$durability <- mydata$dur_gene1 / nTSpY
mydata$durability[mydata$durability > nYears] <- nYears

summary(mydata)
# dim(mydata)
# sum(duplicated(mydata[,2:9]))  ## check if seeds are different for a given set of parameters
# boxplot(audpc_rel_S ~ mutation_prob+fitness_cost+aggreg, data=mydata, ylim=c(0.46, 0.61))


## Aggregation of variables : mean of replicates
outputs <- c("durability", "gla_rel_S", "gla_rel_R", "susceptible_patho2", "resistant_patho2")
names(outputs) <- c("Durability", "S cultivar", "R cultivar", "S cultivar", "R cultivar")
mydata_mean <- aggregate(mydata[,outputs], by=list(aggreg=mydata$aggreg, fitness_cost=mydata$fitness_cost
                                               , mutation_prob=mydata$mutation_prob
                                               , R_efficiency=mydata$R_efficiency
                                               , time_to_activ_exp=mydata$time_to_activ_exp
                                               , target_trait=mydata$target_trait)
                     , FUN=mean)
summary(mydata_mean)
# chart.Correlation(mydata_mean[,outputs], histogram=TRUE, pch=19)


###################################
####         Heat maps         ####
###################################
# subDir <- "heatMaps"
# if (!file.exists(subDir))
#     dir.create(subDir)

COLORS <- cb_palette[c("darkred", "orange", "yellow", "green", "blue")]
COLORS2 <- cb_palette[c("darkred", "darkred", "orange", "yellow", "green", "blue")]
dataToPlot <- melt(mydata_mean, measure.vars=outputs)
levels(dataToPlot$variable) <- outputs
levels(dataToPlot$aggreg) <- c("A) Aggregated landscape", "B) Fragmented landscape")
levels(dataToPlot$fitness_cost) <- c("Fitness cost = 0","Fitness cost = 0.25", "Fitness cost = 0.50")
cultivar_labs <- names(outputs)
names(cultivar_labs) <- outputs

for (tr in traits){ 
    
    plots <- list()
    p=0
    for (a in levels(dataToPlot$aggreg)){
        ## Durability
        p <- p+1
        plots[[p]] <- ggplot(dataToPlot[dataToPlot$target_trait==tr & dataToPlot$aggreg==a & dataToPlot$variable=="durability",]
                             , aes(x=time_to_activ_exp, y=R_efficiency)) + 
            facet_grid(.~fitness_cost) + geom_tile(aes(fill=value)) +
            facet_rep_grid(.~fitness_cost) + 
            # coord_capped_cart(bottom='both', left='both') +
            # coord_fixed() +
            scale_y_continuous(breaks=seq(0,1,.2), labels=paste(seq(0,100,20),"%",sep="")) + 
            scale_x_continuous(breaks=seq(0,120,30)) + 
            xlab("Age of resistance activation (days)") +
            ylab("Resistance efficiency") + 
            ggtitle(paste(a)) +
            theme(aspect.ratio=1, panel.spacing = unit(-1, "lines")
                  , axis.text.x=element_text(angle=40, hjust=1, vjust=1, size=rel(1))
                  , panel.background = element_blank(), axis.line = element_line(colour = "black")
                  , panel.grid.major = element_blank(), panel.grid.minor = element_blank()) +  ## to remove grid
            # guides(fill=FALSE) +    ## to remove the legend
            labs(fill="Durability") + 
            theme(legend.title=element_text(face="bold", size=9)) +
            scale_fill_gradientn(colours=COLORS, limits=c(0,30), breaks=seq(0,30,10), labels=paste(seq(0,30,10)," years",sep=""))
    } ## for a
    
    tiff(paste("heatMap", run, "durab_", tr, ".tiff", sep=""),width=350,height=210,units='mm',compression='lzw',res=300)
    multiplot(plotlist=plots, layout=matrix(1:2,nrow=1,byrow=FALSE))
    dev.off()
    
    plots <- list()
    p=0
    for (a in levels(dataToPlot$aggreg)){
        ## GLA
        p <- p+1
        plots[[p]] <- ggplot(dataToPlot[dataToPlot$target_trait==tr & dataToPlot$aggreg==a & substr(dataToPlot$variable,1,3)=="gla",]
                             , aes(x=time_to_activ_exp, y=R_efficiency)) + 
            facet_grid(variable~fitness_cost) + geom_tile(aes(fill=value)) +
            facet_rep_grid(variable~fitness_cost, labeller=labeller(variable=cultivar_labs)) + 
            # coord_capped_cart(bottom='both', left='both') +
            # coord_fixed() +
            scale_y_continuous(breaks=seq(0,1,.2), labels=paste(seq(0,100,20),"%",sep="")) + 
            scale_x_continuous(breaks=seq(0,120,30)) + 
            xlab("Age of resistance activation (days)") +
            ylab("Resistance efficiency") + 
            ggtitle(paste(a)) +
            theme(aspect.ratio=1, panel.spacing = unit(-1, "lines")
                  , axis.text.x=element_text(angle=40, hjust=1, vjust=1, size=rel(1))
                  , panel.background = element_blank(), axis.line = element_line(colour = "black")
                  , panel.grid.major = element_blank(), panel.grid.minor = element_blank()) +  ## to remove grid
            # guides(fill=FALSE) +    ## to remove the legend
            labs(fill="Disease control\n(GLA)") + 
            theme(legend.title=element_text(face="bold", size=9)) +
            scale_fill_gradientn(colours=COLORS2, limits=c(0.3,1), breaks=seq(0,1,0.1)
                                 , values=rescale(c(0.3, 0.33, 0.38, 0.40, 0.42, 1), from=c(0.3, 1)))

        
        ## CONTRIBUTION
        p <- p+1
        plots[[p]] <- ggplot(dataToPlot[dataToPlot$target_trait==tr & dataToPlot$aggreg==a 
                                        & (substr(dataToPlot$variable,1,3)=="sus" | substr(dataToPlot$variable,1,3)=="res"),]
                             , aes(x=time_to_activ_exp, y=R_efficiency)) + 
            facet_grid(variable~fitness_cost) + geom_tile(aes(fill=value)) +
            facet_rep_grid(variable~fitness_cost, labeller=labeller(variable=cultivar_labs)) + 
            # coord_capped_cart(bottom='both', left='both') +
            # coord_fixed() +
            scale_y_continuous(breaks=seq(0,1,.2), labels=paste(seq(0,100,20),"%",sep="")) + 
            scale_x_continuous(breaks=seq(0,120,30)) + 
            xlab("Age of resistance activation (days)") + ylab("Resistance efficiency") + 
            # ggtitle(paste(a)) +
            theme(aspect.ratio=1, panel.spacing = unit(-1, "lines")
                  , axis.text.x=element_text(angle=40, hjust=1, vjust=1, size=rel(1))
                  , panel.background = element_blank(), axis.line = element_line(colour = "black")
                  , panel.grid.major = element_blank(), panel.grid.minor = element_blank()) +  ## to remove grid
            # guides(fill=FALSE) +    ## to remove the legend
            labs(fill="Frequency of\nthe rb pathogen") + 
            theme(legend.title=element_text(face="bold", size=9)) +
            scale_fill_gradientn(colours=c("white", "black"), limits=c(0,1), breaks=seq(0,1,0.2))

    } ## for a
    tiff(paste("heatMap", run, "control_", tr, ".tiff", sep=""),width=350,height=210,units='mm',compression='lzw',res=300)
    multiplot(plotlist=plots, layout=matrix(1:4,nrow=2,byrow=FALSE))
    dev.off()
    
} ## for tr


###################################
####         Analyses          ####
###################################
source("/home/loup/Dropbox/scriptsLPR/functions_lpr.R")

mydata2 <- mydata[mydata$mutation_prob==1E-4 & mydata$fitness_cost==0.5 & mydata$aggreg==0.05,]

## Durability
## id_landscape is significant but with very small effect (explains nearly nothing of the deviance)
mod1 <- glm(cbind(dur_gene1, 3601-dur_gene1) ~  R_efficiency*time_to_activ_exp + target_trait
            , data=mydata2, family=binomial)
summary(mod1)
hist(mod1$residuals, col="grey")
findR_Dexpl(mod1)
drop1(mod1, test="Chisq")

newdata <- expand.grid(time_to_activ_exp=seq(0,120,1), R_efficiency=seq(0,1,0.01), target_trait=traits)
newdata$pred <- predict(mod1, newdata, type="response")


myplot <- ggplot(newdata, aes(x=time_to_activ_exp, y=R_efficiency)) + 
    facet_grid(~target_trait) + geom_tile(aes(fill=pred)) +
    facet_rep_grid(~target_trait) + coord_capped_cart(bottom='both', left='both') +
    # coord_fixed() +
    scale_y_continuous(breaks=seq(0,1,.2), labels=paste(seq(0,100,20),"%",sep="")) + 
    scale_x_continuous(breaks=seq(0,120,30)) + 
    xlab("Age of resistance activation (days)") + ylab("Resistance efficiency") + 
    # ggtitle(paste("Mut. prob:", m, "   Aggreg:", a)) +
    theme(aspect.ratio=1
          , axis.text.x=element_text(angle=40, hjust=1, vjust=1, size=rel(1))
          , panel.background = element_blank(), axis.line = element_line(colour = "black")
          , panel.grid.major = element_blank(), panel.grid.minor = element_blank()) +  ## to remove grid
    labs(fill="Level of\nevolutionary\nor\nepidemiological\ncontrol") + 
    theme(legend.title=element_text(face="bold", size=9)) +
    # guides(fill=FALSE) +   ## to remove the legend
    scale_fill_gradientn(colours=COLORS, limits=c(0,1), breaks=seq(0,1,.25), labels=paste(seq(0,100,25),"%",sep=""))
plot(myplot)


## AUDPC_S
mod2 <- lm(audpc_rel_S ~  R_efficiency*time_to_activ_exp*target_trait, data=mydata2)
summary(mod2)
hist(mod2$residuals, col="grey")
drop1(mod2, test="Chisq")
```
