## Supplementary material for "Durable resistance or efficient disease control? Adult Plant Resistance (APR) at the heart of the dilemma": Raw data: analyseAPR_3.R

```
######################################################
#########            analyseAPR_3.R          ########
######################################################
#### APR14: two resistance genes: MG + APR
#### APR15: two resistance genes: APR + APR
## landscape : S + R1 + R2, 33%-33%-33%, weak aggregation (0.05), Nyears = 30 years
## evolution: mutation_prob=1E-4, fitness_cost=0.5
## Variation of strategy (4 values), target trait (APR14: 4 values, APR15: 10 combinations)
## Reff  (11 values for 0 to 1), TimetoR (10 values from 0 to 90)
## Stochastic replicates: 5 landscapes, 10 replicates per landscape
## Evolutionary outputs: dur_gene1, dur_gene2.
## Epidemiological outputs: gla_rel_S, gla_rel_R1, gla_rel_R2.
## No need for audpc in whole landscape because audpc_tot = audpc_S + audpc_R1 + audpc_R2


rm(list=ls())
graphics.off()

run = "APR_3"


########  GRAPHICAL PACKAGES  ########
library("ggplot2")     ## for function ggplot
library("lemon")       ## for facet_rep_grid()
library("reshape2")     ## for melt
library("scales")       ## for rescale
library("PerformanceAnalytics")  ## for chart.Correlation
source("/home/loup/Dropbox/scriptsLPR/graphics_lpr.r") 
cb_palette <- c("#000000", "#999999", "#E69F00", "#56B4E9", "#009E73", "#F0E442", "#0072B2", "#D55E00", "#CC79A7", "#b71c1c")
names(cb_palette) <- c("black", "grey", "orange", "cyan", "green", "yellow", "blue", "red", "purple", "darkred")
######################################


######################
####     DATA     ####
######################
setwd(paste("/home/loup/Documents/model_resistance/RES/simul", run, sep=""))

## Load data
allFiles <- list.files(getwd())
filesToOpen <- allFiles[grep("simul_landsepi_2022-07-08", allFiles)]

mydata <- data.frame()
for (i in 1:length(filesToOpen)){
    data_tmp <- read.table(paste(filesToOpen[i], "/myDesign.txt", sep=""), sep=" ", dec=".",header=TRUE)
    data_tmp[,1] <- i
    colnames(data_tmp)[1] <- "rep"
    mydata <- rbind(mydata, data_tmp)
}

## Transform into factors
for (k in c("rep", "id_landscape"))
    mydata[,k] <- factor(mydata[,k])
if (run=="APR_14"){
    traits <- c("IR", "LAT", "PR", "IP")
    names(traits) <- c("Infection rate", "Latent period"
                       , "Sporulation rate", "Sporulation duration")
    mydata$target_traits <- factor(mydata$target_traits, levels=traits)
}else{
    traits <- levels(mydata$target_traits)
}
strategies <- c("MO","MI","RO","PY")
names(strategies) <- c("Mosaic", "Mixture", "Rotation", "Pyramiding")
mydata$strat <- factor(mydata$strat, levels=strategies)
cultivars <- c("susceptibl", "resistant1", "resistant2")
names(cultivars) <- c("A) Susceptible cultivar", "B) Resistant cultivar 1", "C) Resistant cultivar 2")

## Transform durability in % of the maximum possible value (i.e. 30 years) 
nYears <- 30
nTSpY <- 120
mydata$durability1 <- mydata$dur_gene1 / nTSpY
mydata$durability1[mydata$durability1 > nYears] <- nYears
mydata$durability2 <- mydata$dur_gene2 / nTSpY
mydata$durability2[mydata$durability2 > nYears] <- nYears

summary(mydata)
# dim(mydata)
# sum(duplicated(mydata[,2:7]))  ## check if seeds are different for a given set of parameters


## Aggregation of variables : mean of replicates
titles <- expand.grid(cultivar=c("susceptible","resistant1","resistant2"), patho=paste("patho",c(1,3,2,4), sep="")) ## patho2 is rb2, patho3 is rb1
titles$name <- paste(titles$cultivar, titles$patho, sep="_")
outputs <- c("durability1", "durability2", "gla_rel_S", "gla_rel_R1", "gla_rel_R2", titles$name)
names(outputs) <- c("Durability MG", "Durability APR"
                    , "S cultivar", "R1 cultivar", "R2 cultivar"
                    , rep(c("wt", paste("rb",c(1,2,12), sep="")), each=3))

mydata_mean <- aggregate(mydata[,outputs], by=list(R_efficiency=mydata$R_efficiency
                                               , time_to_activ_exp=mydata$time_to_activ_exp
                                               , target_traits=mydata$target_traits
                                               , strat=mydata$strat)
                     , FUN=mean)
summary(mydata_mean)
# chart.Correlation(mydata_mean[,outputs], histogram=TRUE, pch=19)


###################################
####         Heat maps         ####
###################################
# subDir <- "heatMaps"
# if (!file.exists(subDir))
#     dir.create(subDir)

COLORS <- cb_palette[c("darkred", "orange", "yellow", "green", "blue")]
COLORS2 <- cb_palette[c("darkred", "darkred", "orange", "yellow", "green", "blue")]
dataToPlot <- melt(mydata_mean, measure.vars=outputs)
levels(dataToPlot$variable) <- outputs
levels(dataToPlot$strat) <- names(strategies)
cultivar_labs <- names(outputs)
names(cultivar_labs) <- outputs

for (tr in traits){ 
    ## Durability
    myPlot <- ggplot(dataToPlot[dataToPlot$target_traits==tr & substr(dataToPlot$variable,1,3)=="dur",]
                         , aes(x=time_to_activ_exp, y=R_efficiency)) + 
        facet_grid(variable~strat) + geom_tile(aes(fill=value)) +
        facet_rep_grid(variable~strat, labeller=labeller(variable=cultivar_labs)) + 
        # coord_capped_cart(bottom='both', left='both') +
        # coord_fixed() +
        scale_y_continuous(breaks=seq(0,1,.2), labels=paste(seq(0,100,20),"%",sep="")) + 
        scale_x_continuous(breaks=seq(0,90,30)) + 
        xlab("Age of activation of the APR gene (days)") + ylab("APR gene efficiency") + 
        ggtitle(paste("A) Target trait:", tr)) +
        theme(aspect.ratio=1, panel.spacing = unit(-1, "lines")
              , axis.text.x=element_text(angle=40, hjust=1, vjust=1, size=rel(1))
              , panel.background = element_blank(), axis.line = element_line(colour = "black")
              , panel.grid.major = element_blank(), panel.grid.minor = element_blank()) +  ## to remove grid
        labs(fill="Durability") + 
        theme(legend.title=element_text(face="bold", size=9)) +
        # guides(fill=FALSE) +   ## to remove the legend    
        scale_fill_gradientn(colours=COLORS, limits=c(0,30), breaks=seq(0,30,10), labels=paste(seq(0,30,10)," years",sep=""))
    
    tiff(paste("heatMap", run, "durab_", tr, ".tiff", sep=""),width=200,height=100,units='mm',compression='lzw',res=300)
    plot(myPlot)
    dev.off()
    
    ## GLA
    myPlot <- ggplot(dataToPlot[dataToPlot$target_trait==tr & substr(dataToPlot$variable,1,3)=="gla",]
                         , aes(x=time_to_activ_exp, y=R_efficiency)) + 
        facet_grid(variable~strat) + geom_tile(aes(fill=value)) +
        facet_rep_grid(variable~strat, labeller=labeller(variable=cultivar_labs)) + 
        # coord_capped_cart(bottom='both', left='both') +
        # coord_fixed() +
        scale_y_continuous(breaks=seq(0,1,.2), labels=paste(seq(0,100,20),"%",sep="")) + 
        scale_x_continuous(breaks=seq(0,120,30)) + 
        xlab("Age of activation of the APR gene (days)") + ylab("APR gene efficiency") + 
        ggtitle(paste("B) Target trait:", tr)) +
        theme(aspect.ratio=1, panel.spacing = unit(-1, "lines")
              , axis.text.x=element_text(angle=40, hjust=1, vjust=1, size=rel(1))
              , panel.background = element_blank(), axis.line = element_line(colour = "black")
              , panel.grid.major = element_blank(), panel.grid.minor = element_blank()) +  ## to remove grid
        # guides(fill=FALSE) +    ## to remove the legend
        labs(fill="Disease control\n(GLA)") + 
        theme(legend.title=element_text(face="bold", size=9)) +
        # scale_fill_gradientn(colours=COLORS, limits=c(0.3,1), breaks=seq(0,1,0.1)
        #                      , values=rescale(c(0.3, 0.32, 0.38, 0.40, 0.42, 1), from=c(0.3, 1)))
        scale_fill_gradientn(colours=COLORS2, limits=c(0.3,1), breaks=seq(0,1,0.1)
                             , values=rescale(c(0.3, 0.36, 0.43, 0.45, 0.47, 1), from=c(0.3, 1)))
    
    
    tiff(paste("heatMap", run, "control_", tr, ".tiff", sep=""),width=200,height=200,units='mm',compression='lzw',res=300)
    plot(myPlot)
    dev.off() 
    
    ## CONTRIBUTION
    plots <- list()
    p=0

    for (k in 1:length(cultivars)){
        p <- p+1
        plots[[p]] <- ggplot(dataToPlot[dataToPlot$target_trait==tr & substr(dataToPlot$variable,1,10)==cultivars[k],]
                             , aes(x=time_to_activ_exp, y=R_efficiency)) + 
            facet_grid(variable~strat) + geom_tile(aes(fill=value)) +
            facet_rep_grid(variable~strat, labeller=labeller(variable=cultivar_labs)) + 
            # coord_capped_cart(bottom='both', left='both') +
            # coord_fixed() +
            scale_y_continuous(breaks=seq(0,1,.2), labels=paste(seq(0,100,20),"%",sep="")) + 
            scale_x_continuous(breaks=seq(0,120,30)) + 
            xlab("Age of resistance activation (days)") + 
            ggtitle(names(cultivars)[k]) +
            theme(aspect.ratio=1, panel.spacing = unit(-1, "lines")
                  , axis.text.x=element_text(angle=40, hjust=1, vjust=1, size=rel(1))
                  , panel.background = element_blank(), axis.line = element_line(colour = "black")
                  , panel.grid.major = element_blank(), panel.grid.minor = element_blank()) +  ## to remove grid
            guides(fill=FALSE) +    ## to remove the legend
            # labs(fill="Frequency in the\npathogen population") + 
            # theme(legend.title=element_text(face="bold", size=9)) +
            scale_fill_gradientn(colours=c("white", "black"), limits=c(0,1), breaks=seq(0,1,0.2))
        if (k==1){
            plots[[p]] <- plots[[p]] + ylab("Resistance efficiency")
        }else{
            plots[[p]] <- plots[[p]] + ylab("")
        }
    } ## for k
    
    tiff(paste("heatMap", run, "contrib_", tr, ".tiff", sep=""),width=350,height=210,units='mm',compression='lzw',res=300)
    multiplot(plotlist=plots, layout=matrix(1:3,nrow=1,byrow=FALSE))
    dev.off()
} ## for tr

## Legend
dataLegend <- data.frame(x=1:10, y=1:10, value=0)
tiff(paste("heatMap", run, "contrib_legend.tiff", sep=""),width=50,height=50,units='mm',compression='lzw',res=300)
ggplot(dataLegend, aes(x=x, y=y)) + 
    geom_tile(aes(fill=value)) +
    
    theme(axis.line=element_blank(),axis.text.x=element_blank(),
          axis.text.y=element_blank(),axis.ticks=element_blank(),
          axis.title.x=element_blank(), axis.title.y=element_blank(),
          panel.background=element_blank(),panel.border=element_blank(),panel.grid.major=element_blank(),
          panel.grid.minor=element_blank(),plot.background=element_blank()) +
    
    labs(fill="Frequency in the\npathogen population") + 
    theme(legend.title=element_text(face="bold", size=9)) +
    scale_fill_gradientn(colours=c("white", "black"), limits=c(0,1), breaks=seq(0,1,0.2))
dev.off()
```
