## Supplementary material for "Durable resistance or efficient disease control? Adult Plant Resistance (APR) at the heart of the dilemma": Raw data: simul_plan_APR_3.R

```
library("landsepi", lib.loc="/home/lrimbaud/R/library")

## NUMERICAL DESIGN
myDesign <- expand.grid(id_landscape = 1:5
                        , R_efficiency = seq(0,1,0.1)
                        , time_to_activ_exp = seq(0,90,10)
                        , target_traits = c("IR", "LAT", "PR", "IP")
                        , strat = c("MO", "MI", "RO", "PY"))

n <- nrow(myDesign)
myDesign <- cbind(simul = 1:n, seed = sample(n*100, n), myDesign)


## Simulation parameters
simul_params <- createSimulParams()
simul_params <- setTime(simul_params, Nyears = 30, nTSpY = 120)

## Pathogen parameters
basic_patho_param <- loadPathogen("rust")
simul_params <- setPathogen(simul_params, basic_patho_param)     
## Initial conditions 
simul_params <- setInoculum(simul_params, 5E-4)

gene1 <- loadGene(name = "gene1", type = "majorGene")
gene2 <- loadGene(name = "gene2", type = "majorGene")
genes <- data.frame(rbind(gene1, gene2), stringsAsFactors = FALSE)

## SIMULATIONS
for (i in 1:n){  
    print(paste("Running simulation", i, "/", n))
    
    ## seed
    simul_params <- setSeed(simul_params, myDesign$seed[i])
    ## landscape
    landscape <- loadLandscape(myDesign$id_landscape[i])
    simul_params <- setLandscape(simul_params, landscape)
    ## Dispersal
    disp_patho <- loadDispersalPathogen(myDesign$id_landscape[i])
    simul_params <- setDispersalPathogen(simul_params, disp_patho)
    
    ## Cultivars
    cultivar1 <- loadCultivar(name = "Susceptible", type = "growingHost")
    if (myDesign$strat[i]=="PY"){
            cultivar2 <- loadCultivar(name = "Resistant", type = "growingHost")
            cultivars <- data.frame(rbind(cultivar1, cultivar2), stringsAsFactors = FALSE)
    }else{
        cultivar2 <- loadCultivar(name = "Resistant1", type = "growingHost")
        cultivar3 <- loadCultivar(name = "Resistant2", type = "growingHost")
        cultivars <- data.frame(rbind(cultivar1, cultivar2, cultivar3), stringsAsFactors = FALSE)      
    }

    simul_params <- setCultivars(simul_params, cultivars)  
    
    ## Allocate cultivars to croptypes
    if (myDesign$strat[i]=="PY"){
        croptypes <- loadCroptypes(simul_params, names = c("Susceptible crop", "Pyramid"))
        croptypes <- allocateCroptypeCultivars(croptypes, "Susceptible crop", "Susceptible")
        croptypes <- allocateCroptypeCultivars(croptypes, "Pyramid", "Resistant")
    }else if (myDesign$strat[i]=="MI"){
        croptypes <- loadCroptypes(simul_params, names = c("Susceptible crop", "Mixture"))
        croptypes <- allocateCroptypeCultivars(croptypes, "Susceptible crop", "Susceptible")
        croptypes <- allocateCroptypeCultivars(croptypes, "Mixture", c("Resistant1", "Resistant2"))
    }else{
        croptypes <- loadCroptypes(simul_params, names = c("Susceptible crop", "Resistant crop 1", "Resistant crop 2"))
        croptypes <- allocateCroptypeCultivars(croptypes, "Susceptible crop", "Susceptible")
        croptypes <- allocateCroptypeCultivars(croptypes, "Resistant crop 1", "Resistant1")
        croptypes <- allocateCroptypeCultivars(croptypes, "Resistant crop 2", "Resistant2")
    }
    simul_params <- setCroptypes(simul_params, croptypes)
    
    
    ## Genes
    genes$efficiency[2] <- myDesign$R_efficiency[i]  ## update resistance efficiency
    genes$time_to_activ_exp[2] <- myDesign$time_to_activ_exp[i]  ## update time to activation
    genes$time_to_activ_var[2] <- myDesign$time_to_activ_exp[i]  ## update time to activation
    genes$mutation_prob <- 1e-4
    genes$target_trait[2] <- as.character(myDesign$target_traits[i])
    
    if (genes$target_trait[2]=="LAT"){
        genes$efficiency[2] <- genes$efficiency[2] / 2
    }
    
    simul_params <- setGenes(simul_params, genes)
    
    ## Allocate genes to cultivars
    if (myDesign$strat[i]=="PY"){
        simul_params <- allocateCultivarGenes(simul_params, "Resistant", c("gene1", "gene2"))
    }else{
        simul_params <- allocateCultivarGenes(simul_params, "Resistant1", c("gene1"))
        simul_params <- allocateCultivarGenes(simul_params, "Resistant2", c("gene2"))
    }
    
    
    ## Allocate croptypes to landscape  
    if (myDesign$strat[i]=="PY" | myDesign$strat[i]=="MI"){
    simul_params <- allocateLandscapeCroptypes(simul_params,
                                               rotation_period = 0,
                                               rotation_sequence = croptypes$croptypeID,
                                               rotation_realloc = FALSE,
                                               prop = c(1/3, 2/3),
                                               aggreg = 0.05,
                                               graphic = FALSE)
    }else if (myDesign$strat[i]=="RO"){
        simul_params <- allocateLandscapeCroptypes(simul_params,
                                                   rotation_period = 1,
                                                   rotation_sequence = list(c(0,1), c(0,2)),
                                                   rotation_realloc = FALSE,
                                                   prop = c(1/3, 2/3),
                                                   aggreg = 0.05,
                                                   graphic = FALSE)
    }else{
        simul_params <- allocateLandscapeCroptypes(simul_params,
                                                   rotation_period = 0,
                                                   rotation_sequence = croptypes$croptypeID,
                                                   rotation_realloc = FALSE,
                                                   prop = c(1/3, 1/3, 1/3),
                                                   aggreg = 0.05,
                                                   graphic = FALSE) 
    }
    
    ## Outputs
    outputList <- loadOutputs(epid_outputs = c("audpc","audpc_rel","gla","gla_rel","contrib"), evol_outputs = "durability")
    simul_params <- setOutputs(simul_params, outputList)
    
    ## run simulation
    res <- runSimul(simul_params, writeTXT = FALSE, graphic = FALSE)
    
    ## Extract outputs
    contrib <- res$epid_outputs$contrib[,,"Whole simulation"]
    myDesign[i,c("audpc_S")] <- mean(res$epid_outputs$audpc$Susceptible, na.rm=TRUE)
    myDesign[i,c("audpc_rel_S")] <- mean(res$epid_outputs$audpc_rel$Susceptible, na.rm=TRUE)
    myDesign[i,c("gla_S")] <- mean(res$epid_outputs$gla$Susceptible, na.rm=TRUE)
    myDesign[i,c("gla_rel_S")] <- mean(res$epid_outputs$gla_rel$Susceptible, na.rm=TRUE)
    if (myDesign$strat[i]=="PY"){
        myDesign[i,c("audpc_R1")] <- mean(res$epid_outputs$audpc$Resistant, na.rm=TRUE)
        myDesign[i,c("audpc_R2")] <- mean(res$epid_outputs$audpc$Resistant, na.rm=TRUE)
        myDesign[i,c("audpc_rel_R1")] <- mean(res$epid_outputs$audpc_rel$Resistant, na.rm=TRUE)
        myDesign[i,c("audpc_rel_R2")] <- mean(res$epid_outputs$audpc_rel$Resistant, na.rm=TRUE)
        myDesign[i,c("gla_R1")] <- mean(res$epid_outputs$gla$Resistant, na.rm=TRUE)
        myDesign[i,c("gla_R2")] <- mean(res$epid_outputs$gla$Resistant, na.rm=TRUE)
        myDesign[i,c("gla_rel_R1")] <- mean(res$epid_outputs$gla_rel$Resistant, na.rm=TRUE)
        myDesign[i,c("gla_rel_R2")] <- mean(res$epid_outputs$gla_rel$Resistant, na.rm=TRUE)
        ## contrib
        contrib <- contrib[c(1,2,2,3),]
    }else{
        myDesign[i,c("audpc_R1")] <- mean(res$epid_outputs$audpc$Resistant1, na.rm=TRUE)
        myDesign[i,c("audpc_R2")] <- mean(res$epid_outputs$audpc$Resistant2, na.rm=TRUE)
        myDesign[i,c("audpc_rel_R1")] <- mean(res$epid_outputs$audpc_rel$Resistant1, na.rm=TRUE)
        myDesign[i,c("audpc_rel_R2")] <- mean(res$epid_outputs$audpc_rel$Resistant2, na.rm=TRUE)
        myDesign[i,c("gla_R1")] <- mean(res$epid_outputs$gla$Resistant1, na.rm=TRUE)
        myDesign[i,c("gla_R2")] <- mean(res$epid_outputs$gla$Resistant2, na.rm=TRUE)
        myDesign[i,c("gla_rel_R1")] <- mean(res$epid_outputs$gla_rel$Resistant1, na.rm=TRUE)
        myDesign[i,c("gla_rel_R2")] <- mean(res$epid_outputs$gla_rel$Resistant2, na.rm=TRUE)
    }
    myDesign[i,c("audpc_tot")] <- mean(res$epid_outputs$audpc$total)
    myDesign[i,c("audpc_rel_tot")] <- mean(res$epid_outputs$audpc_rel$total, na.rm=TRUE)
    myDesign[i,c("gla_tot")] <- mean(res$epid_outputs$gla$total, na.rm=TRUE)
    myDesign[i,c("gla_rel_tot")] <- mean(res$epid_outputs$gla_rel$total, na.rm=TRUE)

    ## contrib
    titles <- expand.grid(cultivar=c("susceptible","resistant1","resistant2","tot"), patho=paste("patho",1:4, sep=""))
    titles$name <- paste(titles$cultivar, titles$patho, sep="_")
    myDesign[i,titles$name] <- as.numeric(contrib)
    
    ## durability
    myDesign[i,c("dur_gene1","dur_gene2")] <- res$evol_outputs$durability
    
    ## Create myDesign.txt at first simulation and then append
    if (i ==1){
        write.table(myDesign[i,], paste(simul_params@OutputDir, "/myDesign.txt", sep="")
                    , append=FALSE, row.names = FALSE, col.names = TRUE)      
    } else {
        write.table(myDesign[i,], paste(simul_params@OutputDir, "/myDesign.txt", sep="")
                    , append=TRUE, row.names = FALSE, col.names = FALSE)
    }
}
```
